# High resolution maps of chromatin reorganization through mouse meiosis reveal novel features of the 3D meiotic structure

**DOI:** 10.1101/2024.03.25.586627

**Authors:** Gang Cheng, Florencia Pratto, Kevin Brick, Xin Li, Benjamin Alleva, Mini Huang, Gabriel Lam, R. Daniel Camerini-Otero

**Affiliations:** Genetics and Biochemistry Branch, NIDDK, National Institutes of Health, Bethesda, MD, USA; KariusDX; Sun Yat-Sen University, School of Medicine, Shen Zhen, China; RNA Regulation Section, NIA, National Institutes of Health, Baltimore, MD, USA

## Abstract

When germ cells transition from the mitotic cycle into meiotic prophase I (MPI), chromosomes condense into an array of chromatin loops that are required to promote homolog pairing and genetic recombination. To identify the changes in chromosomal conformation, we isolated nuclei on a trajectory from spermatogonia to the end of MPI. At each stage along this trajectory, we built genomic interaction maps with the highest temporal and spatial resolution to date. The changes in chromatin folding coincided with a concurrent decline in mitotic cohesion and a rise in meiotic cohesin complexes. We found that the stereotypical large-scale A and B compartmentalization was lost during meiotic prophase I alongside the loss of topological associating domains (TADs). Still, local subcompartments were detected and maintained throughout meiosis. The enhanced Micro-C resolution revealed that, despite the loss of TADs, higher frequency contact sites between two loci were detectable during meiotic prophase I coinciding with CTCF bound sites. The pattern of interactions around these CTCF sites with their neighboring loci showed that CTCF sites were often anchoring the meiotic loops. Additionally, the localization of CTCF to the meiotic axes indicated that these anchors were at the base of loops. Strikingly, even in the face of the dramatic reconfiguration of interphase chromatin into a condensed loop-array, the interactions between regulatory elements remained well preserved. This establishes a potential mechanism for how the meiotic chromatin maintains active transcription within a highly structured genome. In summary, the high temporal and spatial resolution of these data revealed previously unappreciated aspects of mammalian meiotic chromatin organization.

## Introduction

The spatial organization of genomes plays a pivotal role in biological functions (Dekker 2008; Yu Zhang et al. 2012; Marchal, Sima, and Gilbert 2019). The organization of the mammalian genome is hierarchical. Chromatin structures are spatially organized into distinct scales ranging from chromosome territories to compartments, topologically associating domains (TADs), and chromatin loops (Szabo, Bantignies, and Cavalli 2019).

During meiosis, chromosomes undergo dramatic structural changes needed for the proper alignment and segregation of homologs (Zickler and Kleckner 2023). As cells enter meiosis, chromosomes are organized as linear loop arrays, emanating from a proteinaceous axis which include cohesins containing meiotic-specific subunits and axial core elements. Such reorganization leads to linearized chromosomes that resemble the morphology of the mitotic chromosomes (Kleckner, Zickler, and Witz 2013; Zickler and Kleckner 2023). Importantly, and differently from mitosis, this specialized chromosome structure retains active transcription (Ur and Corbett 2021).

Until recently, most of our understanding of meiotic chromatin structure came from cytological studies (Grey and de Massy 2021; Zickler and Kleckner 2023), but technological advances have enabled the interrogation of meiotic chromosome organization through Chromosome Conformation Capture (3C)-based methods (Lieberman-Aiden et al. 2009; Dekker et al. 2002). Recent studies have used Hi-C to detect genome-wide chromatin interactions during mammalian spermatogenesis. Studies both in rhesus monkey and mouse have shown that TADs break down as cells enter meiotic prophase I (Patel et al. 2019; Luo et al. 2020; Zuo et al. 2021; Alavattam et al. 2019; Vara et al. 2019; He et al. 2023). Alongside this attenuation of TADs was a progressive increase of loop size from ∼500 kb to a maximum of 1.8Mb in late meiotic prophase I (Zuo et al. 2021). In yeast, loops were found to be prominently positioned and associated with the chromosome axis (Schalbetter et al. 2019; Grey and de Massy 2021), however, this type of reproducible positioning of loops has not been detected in mammalian spermatocytes. Importantly, the earliest stage assessed in mammalian meiotic cells was meiotic S phase, where loop size is already longer than in mitotic cells (Zuo et al. 2021). The technical difficulties in isolating stages preceding meiotic S has prevented studies investigating chromatin reorganization leading up to meiotic prophase I. Moreover, the chromosomal interactions at fine scales were not examined in previous studies. These types of analyses can be conducted using Micro-C, a derivative of Hi-C, where the genome is fragmented by micrococcal nuclease, instead of restriction enzymes, providing a higher-resolution view of chromosomal interactions (Krietenstein et al. 2020; Hsieh et al. 2020, 2015).

In this study, we used a unique nuclei sorting strategy (Lam et al. 2019) to isolate germ cells at stages ranging from spermatogonia, through the mitotic-to-meiotic transition stages, and up to the end of meiotic prophase I. By performing in-situ Hi-C (Rao et al. 2014) and Micro-C, we explored the trajectory of chromatin reorganization at the highest possible temporal and spatial resolution. We focused on delineating the earliest stage where chromatin becomes “meiotic” and revealed novel features of underlying chromatin organization in meiotic prophase I.

### Isolation of stage-specific nuclei through spermatogenesis

To identify temporal chromatin structure changes during spermatogenesis, we isolated pure populations of germline nuclei using fluorescence-activated nuclei sorting (Figure 1A; S1A; (Lam et al. 2019)). Nuclei from the germ cells that preceded meiotic entry were identified using a sorting paradigm that relies on DNA content and a combination of intranuclear proteins associated with each stage (Figure S1A). Undifferentiated spermatogonia (unDiff.SGA) were identified as 2C nuclei expressing the PLZF (promyelocytic leukemia zinc finger) protein (Buaas et al. 2004; Costoya et al. 2004) (Figure S1A-C). Other “pre-meiotic” populations up to meiotic G1 were identified using a combination of DMRT1, which suppresses meiotic entry and is expressed throughout spermatogonial development (Matson et al. 2010), and STRA8, which can trigger meiotic entry but is expressed both in spermatogonia (Zhou et al. 2008; Endo et al. 2015) and in cells that enter MPI (Endo et al. 2015) (Figure S1A-C). Meiotic S phase, Leptotene, Zygotene, Pachytene, and Diplotene nuclei were isolated as described previously (Lam et al. 2019; Pratto et al. 2021). This nuclei sorting approach was shown to generate populations by up to 94% purity (Lam et al. 2019).

**Figure 1.**
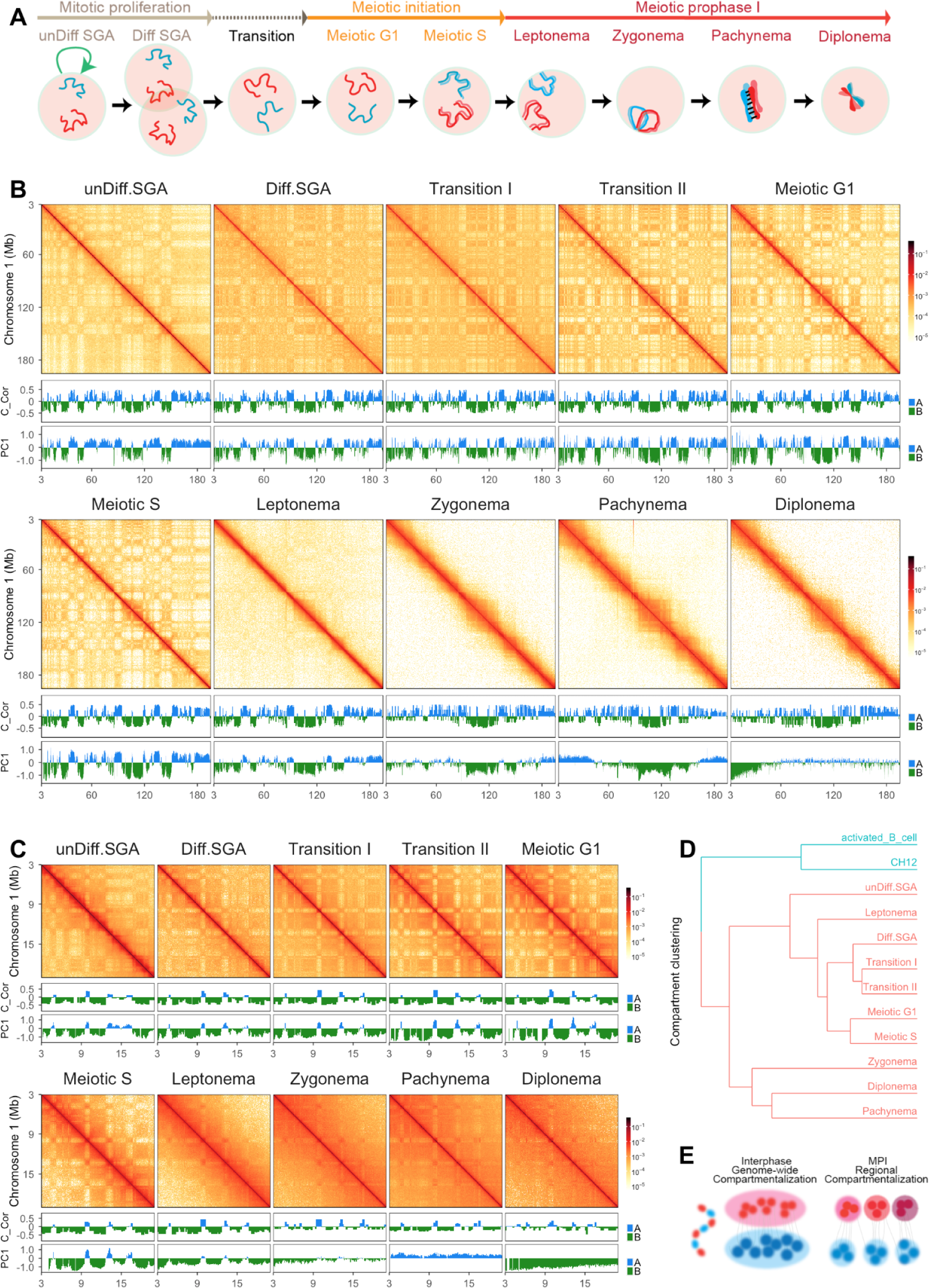
Genome-wide chromatin reorganization throughout spermatogenesis. (A) A schematic of cell populations examined in this study. (B) Genome-wide chromatin reorganization of chromosome 1 derived from Hi-C data. Top: Matrices of chromosome 1 at a resolution of 100 Kb. Middle: Track shows the compartment scores generated by Calder (Y. Liu et al. 2021). To compare with the eigenvector values, we subtracted 0.5 to each score. Bottom: Track shows the eigenvector values for the first principal component of the Hi-C matrix calculated at a resolution of 100 Kb. Blue: active regions; Green: inactive regions. Calder and PC1 generated comparable profiles, with the exception of the zygonema, pachynema, and diplonema stages. (C) A snapshot of Hi-C matrices at chromosome 1: 3 - 20 Mb. Top: Matrices were plotted at a resolution of 50 Kb. Middle: Track shows the compartment scores generated by Calder (Y. Liu et al. 2021). To compare with the eigenvector values, we subtracted 0.5 to each score. Bottom: Track shows the eigenvector values for the first principal component of the Hi-C matrix calculated at a resolution of 100 Kb. Blue: active regions; Green: inactive regions. Both Calder and PC1 were able to differentiate compartment intervals by the leptotene stage. In the zygonema, pachynema and diplonema stages, where only local “checker-board”-like was observed, Calder maintains the ability to identify compartment intervals whereas PC1 was unable to distinguish the genomic regions with visible differences in contact preference. (D) Clustering of all stages based on compartmental annotation derived from Calder (Y. Liu et al. 2021). (E) A schematic of the local compartmentalization during MPI. The schematic was adapted from (Hildebrand and Dekker 2020). Left: Blue and red represent alternating A (active) and B (inactive) compartment intervals on a chromatin fiber. Middle: Genome-wide compartmentalization into two big domains. Right: Compartmentalization is restrained to local, forming small domains.

We performed in-situ Hi-C on all stages of isolated nuclei. All replicates were pooled after confirming their reproducibility (Figure S2). To examine the interactions at high resolution, we employed Micro-C (Figure S3; Hsieh et al. 2015; Krietenstein et al. 2020; Hsieh et al. 2020)), on a subset of nuclei isolated from Meiotic S through diplonema and generated contact maps containing up to 3 billion interactions (Table 1), a number of contacts that would provide an average of 10 unique contacts per 10 kb bin-pairs and that was deemed enough to identify most of detectable loops (Harris et al. 2023).

**Table 1:**
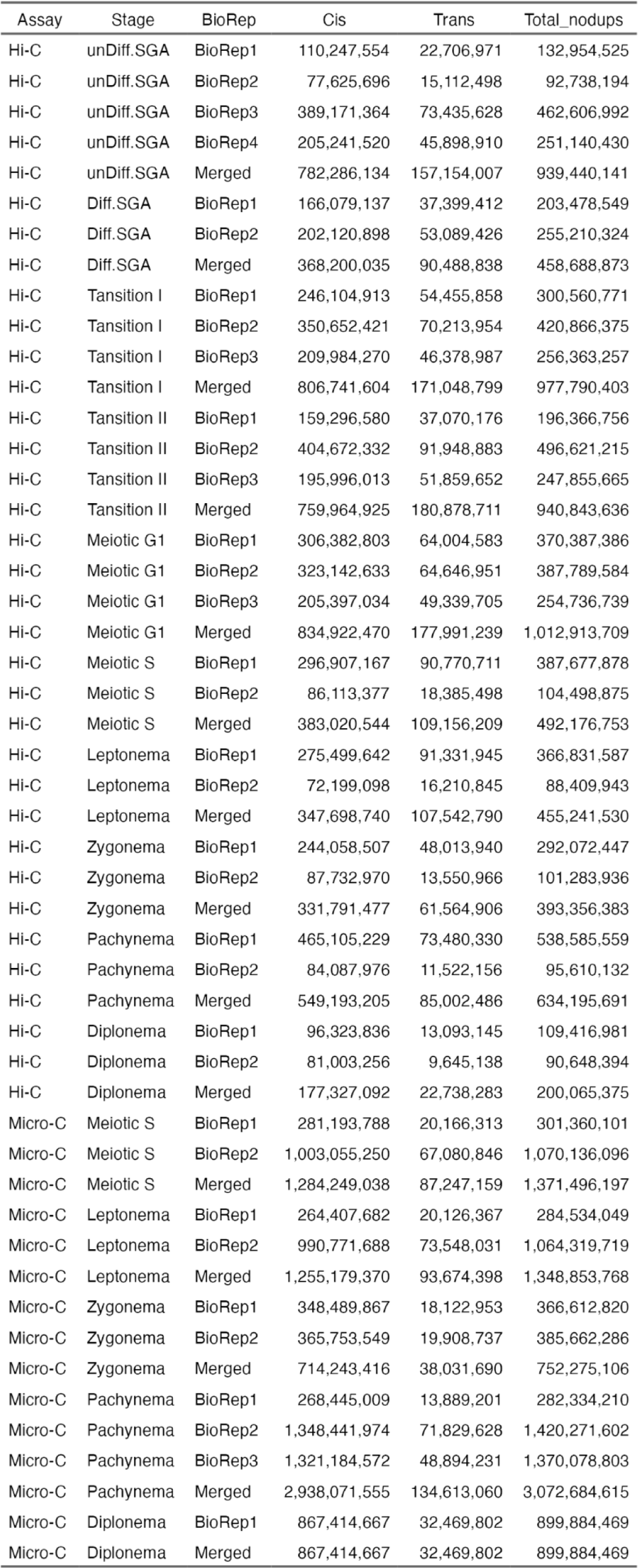
Hi-C & Micro-C library summary.

| Table1: Hi-C & Micro-C library summary |  |  |  |  |  |
| --- | --- | --- | --- | --- | --- |
| Assay | Stage | BioRep | Cis | Trans | Total_nodups |
| Hi-C | unDiff.SGA | BioRep1 | 110,247,554 | 22,706,971 | 132,954,525 |
| Hi-C | unDiff.SGA | BioRep2 | 77,625,696 | 15,112,498 | 92,738,194 |
| Hi-C | unDiff.SGA | BioRep3 | 389,171,364 | 73,435,628 | 462,606,992 |
| Hi-C | unDiff.SGA | BioRep4 | 205,241,520 | 45,898,910 | 251,140,430 |
| Hi-C | unDiff.SGA | Merged | 782,286,134 | 157,154,007 | 939,440,141 |
| Hi-C | Diff.SGA | BioRep1 | 166,079,137 | 37,399,412 | 203,478,549 |
| Hi-C | Diff.SGA | BioRep2 | 202,120,898 | 53,089,426 | 255,210,324 |
| Hi-C | Diff.SGA | Merged | 368,200,035 | 90,488,838 | 458,688,873 |
| Hi-C | Tansition I | BioRep1 | 246,104,913 | 54,455,858 | 300,560,771 |
| Hi-C | Tansition I | BioRep2 | 350,652,421 | 70,213,954 | 420,866,375 |
| Hi-C | Tansition I | BioRep3 | 209,984,270 | 46,378,987 | 256,363,257 |
| Hi-C | Tansition I | Merged | 806,741,604 | 171,048,799 | 977,790,403 |
| Hi-C | Tansition II | BioRep1 | 159,296,580 | 37,070,176 | 196,366,756 |
| Hi-C | Tansition II | BioRep2 | 404,672,332 | 91,948,883 | 496,621,215 |
| Hi-C | Tansition II | BioRep3 | 195,996,013 | 51,859,652 | 247,855,665 |
| Hi-C | Tansition II | Merged | 759,964,925 | 180,878,711 | 940,843,636 |
| Hi-C | Meiotic G1 | BioRep1 | 306,382,803 | 64,004,583 | 370,387,386 |
| Hi-C | Meiotic G1 | BioRep2 | 323,142,633 | 64,646,951 | 387,789,584 |
| Hi-C | Meiotic G1 | BioRep3 | 205,397,034 | 49,339,705 | 254,736,739 |
| Hi-C | Meiotic G1 | Merged | 834,922,470 | 177,991,239 | 1,012,913,709 |
| Hi-C | Meiotic S | BioRep1 | 296,907,167 | 90,770,711 | 387,677,878 |
| Hi-C | Meiotic S | BioRep2 | 86,113,377 | 18,385,498 | 104,498,875 |
| Hi-C | Meiotic S | Merged | 383,020,544 | 109,156,209 | 492,176,753 |
| Hi-C | Leptonema | BioRep1 | 275,499,642 | 91,331,945 | 366,831,587 |
| Hi-C | Leptonema | BioRep2 | 72,199,098 | 16,210,845 | 88,409,943 |
| Hi-C | Leptonema | Merged | 347,698,740 | 107,542,790 | 455,241,530 |
| Hi-C | Zygonema | BioRep1 | 244,058,507 | 48,013,940 | 292,072,447 |
| Hi-C | Zygonema | BioRep2 | 87,732,970 | 13,550,966 | 101,283,936 |
| Hi-C | Zygonema | Merged | 331,791,477 | 61,564,906 | 393,356,383 |
| Hi-C | Pachynema | BioRep1 | 465,105,229 | 73,480,330 | 538,585,559 |
| Hi-C | Pachynema | BioRep2 | 84,087,976 | 11,522,156 | 95,610,132 |
| Hi-C | Pachynema | Merged | 549,193,205 | 85,002,486 | 634,195,691 |
| Hi-C | Diplonema | BioRep1 | 96,323,836 | 13,093,145 | 109,416,981 |
| Hi-C | Diplonema | BioRep2 | 81,003,256 | 9,645,138 | 90,648,394 |
| Hi-C | Diplonema | Merged | 177,327,092 | 22,738,283 | 200,065,375 |
| Micro-C | Meiotic S | BioRep1 | 281,193,788 | 20,166,313 | 301,360,101 |
| Micro-C | Meiotic S | BioRep2 | 1,003,055,250 | 67,080,846 | 1,070,136,096 |
| Micro-C | Meiotic S | Merged | 1,284,249,038 | 87,247,159 | 1,371,496,197 |
| Micro-C | Leptonema | BioRep1 | 264,407,682 | 20,126,367 | 284,534,049 |
| Micro-C | Leptonema | BioRep2 | 990,771,688 | 73,548,031 | 1,064,319,719 |
| Micro-C | Leptonema | Merged | 1,255,179,370 | 93,674,398 | 1,348,853,768 |
| Micro-C | Zygonema | BioRep1 | 348,489,867 | 18,122,953 | 366,612,820 |
| Micro-C | Zygonema | BioRep2 | 365,753,549 | 19,908,737 | 385,662,286 |
| Micro-C | Zygonema | Merged | 714,243,416 | 38,031,690 | 752,275,106 |
| Micro-C | Pachynema | BioRep1 | 268,445,009 | 13,889,201 | 282,334,210 |
| Micro-C | Pachynema | BioRep2 | 1,348,441,974 | 71,829,628 | 1,420,271,602 |
| Micro-C | Pachynema | BioRep3 | 1,321,184,572 | 48,894,231 | 1,370,078,803 |
| Micro-C | Pachynema | Merged | 2,938,071,555 | 134,613,060 | 3,072,684,615 |
| Micro-C | Diplonema | BioRep1 | 867,414,667 | 32,469,802 | 899,884,469 |
| Micro-C | Diplonema | Merged | 867,414,667 | 32,469,802 | 899,884,469 |

### Local compartmentalization of meiotic prophase I chromosomes is preserved despite the loss of higher-order structures

Previous studies have shown that in interphase, the inherent affinity within active (A) and inactive (B) genomic regions leads to the spatial partitioning of chromosomes into two distinct compartments, which is exhibited as a genome-wide “checker-board”-like pattern in Hi-C maps (Lieberman-Aiden et al. 2009). In meiosis, most studies, excluding Vara et al, have reported that the typical A and B compartments are still observed (Table 2; Grey and de Massy 2021; Ur and Corbett 2021; Patel et al. 2019; Luo et al. 2020; Zuo et al. 2021b; Alavattam et al. 2019; Vara et al. 2019; He et al. 2023). Given the reorganization of meiotic chromosomes into axis-loop arrays, these observations were unexpected. Therefore, we revisited this question by leveraging the high resolution of our data.

**Table 2:**
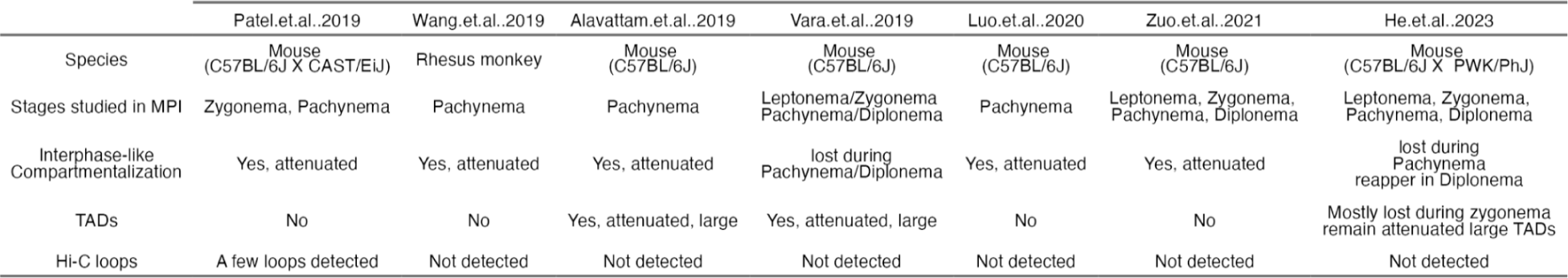
Summary of chromatin conformation studied by Hi-C during mammalian spermatogenesis.

|  |  |  |  |  |  |  |  |
| --- | --- | --- | --- | --- | --- | --- | --- |
|  | Patel.et.al..2019 | Wang.et.al..2019 | Alavattam.et.al..2019 | Vara.et.al..2019 | Luo.et.al..2020 | Zuo.et.al..2021 | He.et.al..2023 |
| Species | Mouse<br>(C57BL/6J X CAST/EiJ) | Rhesus monkey | Mouse<br>(C57BL/6J) | Mouse<br>(C57BL/6J) | Mouse<br>(C57BL/6J) | Mouse<br>(C57BL/6J) | Mouse<br>(C57BL/6J X PWK/PhJ) |
| Stages studied in MPI | Zygonema, Pachynema | Pachynema | Pachynema | Leptonema/Zygonema<br>Pachynema/Diplonema | Pachynema | Leptonema, Zygonema,<br>Pachynema, Diplonema | Leptonema, Zygonema,<br>Pachynema, Diplonema |
| Interphase-like<br>Compartmentalization | Yes, attenuated | Yes, attenuated | Yes, attenuated | lost during<br>Pachynema/Diplonema | Yes, attenuated | Yes, attenuated | lost during<br>Pachynema<br>reappear in Diplonema |
| TADs | No | No | Yes, attenuated, large | Yes, attenuated, large | No | No | Mostly lost during zygonema<br>remain attenuated large TADs |
| Hi-C loops | A few loops detected | Not detected | Not detected | Not detected | Not detected | Not detected | Not detected |

We found that the whole-chromosome “checker-board”-like pattern was present in germline interphase cells and persisted up to when cells entered the meiotic S phase (Figure 1B). The pattern of far-cis interactions began to diminish immediately as cells entered leptonema (Figure 1B; Figure S4A; S5) and vanished entirely at pachynema (Figure 1B; Figure S4A; S5). Using principal component analysis (PCA), we were able to assign A/B compartment intervals along the genome from undifferentiated spermatogonia to leptonema (Figure 1B,C). However, A/B compartment differentiation by PCA analysis became less distinct in zygonema and was completely lost in pachynema and diplonema (Figure 1B; Figure S4A). These observations were consistent across all replicates in both our Hi-C and Micro-C data (Figure S6). To explore the differences in A/B compartment detection (Table 2), we first added meiotic S interaction data into pachytene interaction data and found that even 5% *in silico* “contamination” resulted in the detection of typical A/B compartmentalization by PCA in pachynema (Figure S7). Second, most previous studies performed cell sorting without fixation. Prolonged *in vitro* manipulation of live spermatocytes potentially slightly altered the intrinsic properties of cells. In contrast, our data were derived from cells that were fixed immediately after dissection, potentially ensuring better preservation of chromatin structure.

As observed before (Wang et al. 2019a; He et al. 2023), despite the loss of A/B compartments, we observed a local “checker-board”-like pattern that was evident across all stages. We utilized Calder (Y. Liu et al. 2021), an algorithm designed to use short-range intra-chromosomal interactions to classify domains (Figure 1B,C; Figure S4A,B). These local compartments were found across all stages, from Hi-C or Micro-C data, even though the strength of interactions varied (Figure 1C; Figure S4B).

Clustering analyses revealed that all germ cells were grouped together, and were clearly distinct from the cluster with CH12 and activated B cells, an example of somatic interphase cells. Within the germ cell group, late meiotic prophase I stages (zygotene, pachytene and diplotene) are distinct from spermatogonia and early prophase stages. The local compartments are aligned with transcriptional units and likely to be correlated with the transcriptional hubs described in (Patel et al. 2019). Distinct from previous reports (He et al. 2023), we found that these fine-scale compartments are not exclusive to the meiotic prophase I chromatin and indicated that compartment intervals maintain their shorter-distance contact preference throughout meiotic prophase I. However, the restraint of long-distance interactions, potentially due to the axis-loop array organization, prevents the whole chromosome from segregating into large compartments. Consequently, only domains of smaller size are being preserved (Figure 1E).

### TAD structures change prior to meiotic entry

Meiosis is characterized by the reconfiguration of chromosomes into condensed loop arrays upon entering into meiotic prophase I. To define the transformation from interphase chromatin folding to meiotic loops, we examined the changes in chromosomal folding during mitotic-to-meiotic transition at the scale of TADs. These are typical interphase structures where interactions occur within domains rarely extending to regions beyond their boundaries (Figure 2A; Beagan and Phillips-Cremins 2020). We observed noticeable changes in TAD structures prior to meiotic entry (Figure 2B). While the undifferentiated spermatogonia (unDiff.SGA) interaction map displayed typical TAD structures, an attenuation was evident already in differentiating spermatogonia (Diff.SGA). TADs were further weakened upon initiation of meiosis (Meiotic G1/S), becoming barely visible by leptonema (Figure 2B).

**Figure 2.**
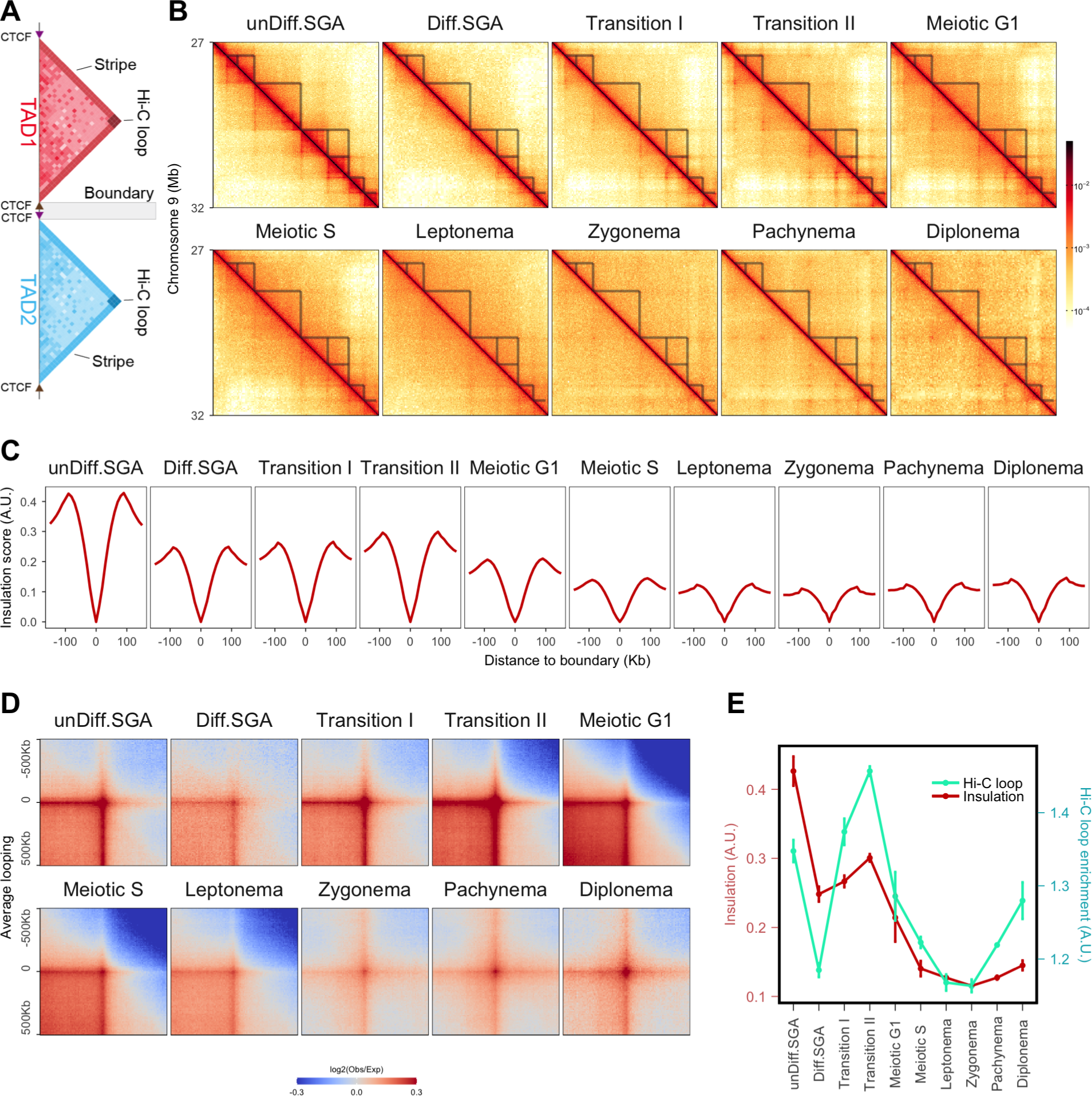
Chromatin folding changes during the mitotic-to-meiotic transition. (A) A schematic of typical TAD structure in Hi-C matrix. Interactions are enriched within TAD1 and TAD2; and isolated between TAD1 and TAD2. CTCF binding sites with forward and backward oriented motifs are indicated by arrows. (B) Hi-C matrices of chromosome 9 at a resolution of 10 Kb (region: 27 Mb to 32 Mb). TADs were located from unDiff.SGA and annotated as tranengles in all stages. (C) Average insulation scores flanking 150 Kb of conserved TAD boundaries. The conserved TAD boundaries were defined by overlapping the boundaries identified from unDiff.SGA and mES cells. The insulation scores were called at a resolution of 10 Kb with a window size of 100 Kb; the minima were normalized to zero. (D) Aggregated interactions between CTCF binding sites with convergent motifs. Each map was plotted at a resolution of 10 Kb, with a flanking region of 500 Kb. (E) Quantitative summary of Figure 2B and 2C. Insulation was calculated as the difference between the maxima and minima values. Mean values of biological replicates were calculated (mean±sd); Hi-C loop enrichment was calculated as the mean of the 3 central pixels in the heatmaps. The mean values of enrichment from different biological replicates were calculated (mean±sd).

Insulation profiles are used to detect TADs and to quantify the difference in average contacts inside and outside the domain (Crane et al. 2015). We detected TADs in unDiff.SGA and kept the boundaries that overlap those detected in mouse embryonic stem cells (mES) (Yan et al. 2018) to derive a highly conserved set of TAD boundaries. We then plotted the insulation score at these sites across all stages. Consistent with our observations from Hi-C matrices, insulation between TADs in Diff.SGA was reduced to roughly half of that in unDiff.SGA (Figure 2C,E). Insulation scores increased slightly during Transition stages, then significantly decreased after meiosis initiation (Figure 2C,E).

Another prominent feature of TADs is the intense point-to-point interactions at their corners termed “Hi-C” loops (Gassler et al. 2017; Figure 2A). In interphase cells, Hi-C loops predominantly coincide with convergent CTCF binding sites (Sanborn et al. 2015; Rao et al. 2014; Fudenberg et al. 2017; Y. Li et al. 2020). To compare Hi-C loop strength among stages, we aggregated interactions between all pairs of convergent CTCF binding sites (Figure 2D). As expected, changes in Hi-C loop strength correlated with insulation changes in most stages (Figure 2D,E). However, an exception was observed at the Transition stages (Figure 2D,E). While both Transition stages displayed lower insulation than unDiff.SGA, their Hi-C loops were markedly stronger (Figure 2E). These data show that germ cells change their chromatin folding prior to entering prophase I, potentially as a preparatory step for meiotic loop formation. Our high temporal resolution allowed us to pinpoint the specific stage at which interphase chromatin structure begins to transition into meiotic chromatin.

### Extended loop formation is associated with the coordinated changes in mitotic and meiotic cohesins during the mitotic-to-meiotic transition

We sought to identify the mechanism that drives the changes in chromatin conformation during mitotic-to-meiotic transition. Properties of chromosome folding can be assessed by calculating the contact probability as a function of genomic distance *P(s) (Gassler et al. 2017).* We observe a gradual change in the curves as they progress from undifferentiated spermatogonia into meiosis. The *P(s)* profile of unDiff.SGA exhibited a typical “hump” at a contact distance less than 1 Mb (Figure 3A, gray area). The “hump” became less pronounced as cells progressed to the Diff.SGA and the Transition I stage (Figure 3A). Subsequently, a new “hump”, beyond 1 Mb, emerged during the Transition II stage and onward (Figure 3A). A *P(s)* curve derived from cells defective in cohesin loading and/or lacking loop extrusion activities is shown as a reference (gray line).

**Figure 3.**
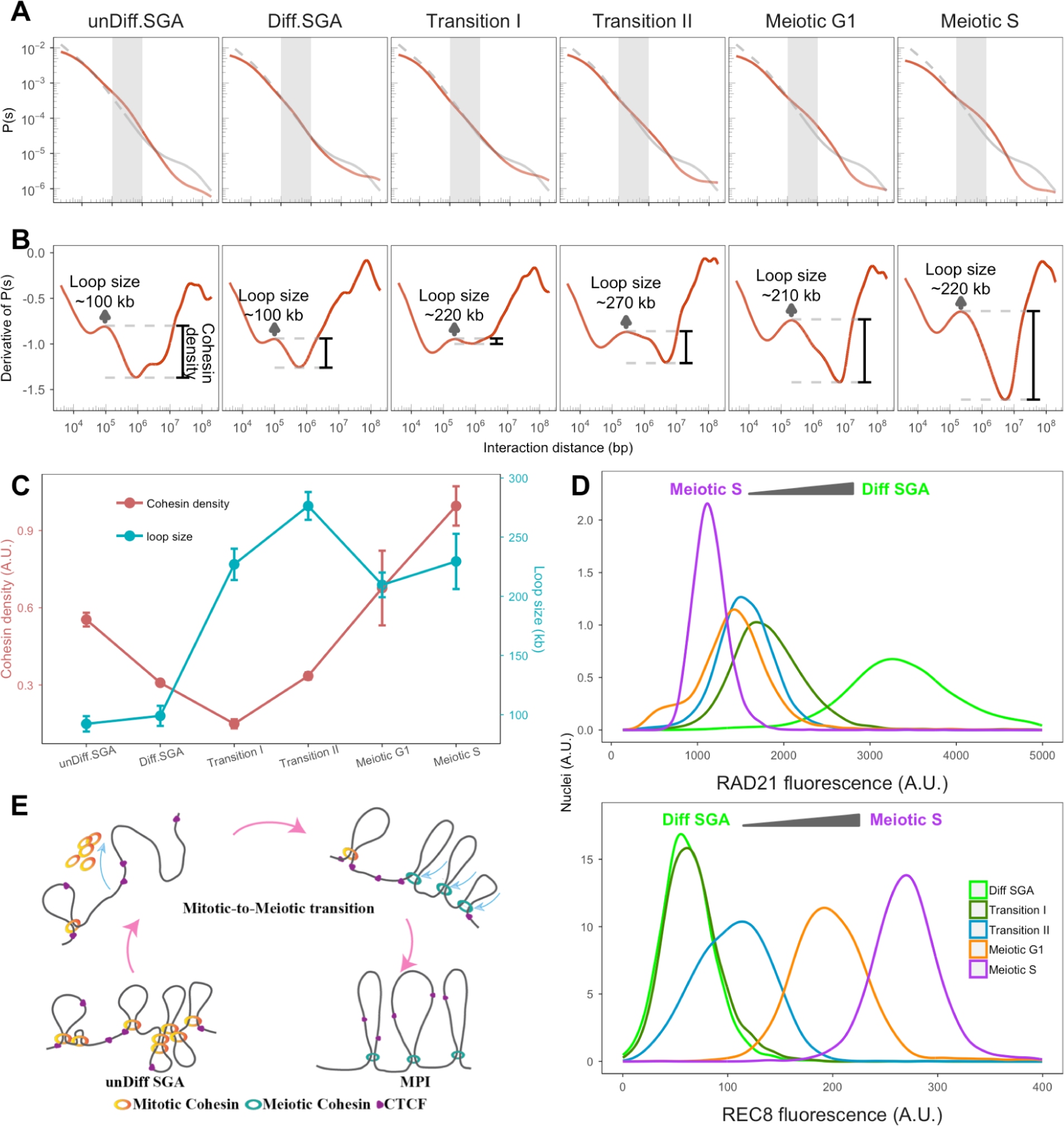
Chromatin reorganization is associated with changes in cohesin during the mitotic-to-meiotic transition. (A) The contact probabilities are plotted as a function of interaction distance P(s). The dashed gray lines are contact probabilities from samples without NIPBL (Schwarzer et al. 2017). The gray regions indicate the scale of TADs. (B) The derivatives of their corresponding P(s) curves. The first local maximums are indicated as arrows, designating the estimated average loop size. The depth of the valley is indicated by the black bar (cohesin density). (C) Quantitative summary of Figure 3B. The relative cohesin density was determined by subtracting the derivative value of the first “bump” and the minimum value. The mean value of biological replicates was calculated and the standard deviation is shown as error bar (mean±*sd*); loop size is the contact distance of the first “bump”. The mean loop size was calculated and the standard deviation is shown as an error bar. (D) The mitotic and meiotic cohesin abundance at “pre-meiotic” stages. Top: The profile of RAD21 fluorescence detected by flow cytometry from Diff.SGA to Meiotic S. Bottom: The profile of REC8 fluorescence detected by flow cytometry from Diff SGA to Meiotic S. (E) A schematic of the proposed model for chromatin folding changes during the mitotic-to-meiotic transition.

Previous work using polymer simulations has shown that the derivative of *P(s)* can reveal the average loop lengths and the density of loops on the chromosomes (Gassler et al. 2017). As indicated by the first local maxima of the derivative, unDiff.SGA have loops with an average size of 100 Kb (Figure 3B,C), which is similar to that found in most somatic interphase cells (Akgol Oksuz et al. 2021; Gassler et al. 2017; Abramo et al., n.d.). The average loop size remained largely constant as cells transitioned to Diff.SGA (Figure 3B,C). However, the “valley” following the first “bump” became shallower in Diff.SGA (Figure 3B); and the derivative curve was nearly flat upon entering into the Transition I stage. According to the polymer simulations (Gassler et al. 2017) this would signify a decrease in cohesin loading and/or loop extrusion activity (Figure 3B). In addition, the remaining loops were more than two times larger than those of the previous stage, averaging around 220 Kb (Figure 3B,C). As cells progressed into meiosis, the average loop size persisted above 200 Kb, with cohesin density increasing until the meiotic S stage (Figure 3B,C). After cells entered meiotic prophase I, loop size kept increasing, reaching the maximum at pachynema (Figure S5A,B and (Patel et al. 2019)). Contrary to previous work (Zuo et al. 2021a), our data indicated a decrease in loop size during diplonema (Figure S5A,B).

Interestingly, transition stage I derivative curves mirrored the pattern seen during telophase when somatic cells exit mitosis (Abramo et al., n.d.). During telophase, there is a condensin-to-cohesin transition, where condensins are evicted from the chromosomes before cohesin binding, leading to an intermediate state of chromatin folding (Abramo et al., n.d.). Similarly, during entry to meiosis, cohesin’s kleisin transitions from RAD21 to meiotic specific kleisins. Altogether, this suggests that cohesin dynamics are disrupted in the transition stages.

To investigate this hypothesis, we carried out flow cytometry analyses to track the changes in mitotic and meiotic cohesin subunits of protein levels across different stages. In line with our hypothesis, RAD21, the α-kleisin of the mitotic cohesin complex, declined immediately as cells entered Transition I, and remained at a low level in subsequent stages (Figure 3D; Figure S9A). Conversely, REC8, a meiotic-specific α-kleisin, began to increase at Transition II, peaking at Meiotic S (Figure 3D; Figure S9B). The shifts in protein levels of RAD21 and REC8 was further confirmed by Western blot (Figure S10). A rise in RAD21L, another meiotic-specific α-kleisin, was not observed until Meiotic S (Figure S9C). This shows that REC8 is the meiotic-specific α-kleisin involved with the chromatin structural changes observed during the mitotic-to-meiotic transition. At the same time, no significant changes in CTCF levels were observed (Figure S9D).

Our data unveiled a crucial temporal point that delineates the shift from mitotic cohesins to meiotic cohesins. The data indicates that when germ cells begin to enter meiosis, mitotic cohesins are unloaded from chromosomes before meiotic cohesins are loaded, leaving the chromosomes largely devoid of loops at this early stage.

### CTCF stabilizes the bound loci at the base of the meiotic loop array

As shown above and consistent with previous studies, we observed a progressive reduction in interaction enrichment within TADs throughout meiotic prophase I (Table 2; Patel et al. 2019; Alavattam et al. 2019; Wang et al. 2019b; Vara et al. 2019; Zuo et al. 2021b); no TADs were discernible even from the pachytene Micro-C interaction matrix built from 3 billion contacts (Figure 4A). Interestingly, and unlike previous studies where no reproducible loop locations were detected (Figure S11A), a punctuate grid-like pattern was observed throughout meiotic prophase I (Figure 2A, Figure S11A,B,C). We initially focused on our Micro-C data on pachynema, since this deeply-sequenced data provided the best visibility and allowed us to perform a fair comparison with publicly available mES Micro-C data (3.3 billion contacts; Hsieh et al. 2020). In the matrix, dots indicating higher frequency of interactions between two loci, were clearly visible, with anchors coinciding with testes CTCF ChIP-seq peaks (Figure 4A). In interphase cells, the interplay between CTCF and cohesins shape architectural features that are detected by Hi-C. Cohesin loops are most frequently found between pairs of convergent CTCF sites and they mediate the formation of TADs (Fudenberg et al. 2017). During mitosis, CTCF is evicted from chromosomes (Gibcus et al. 2018), but there is abundant evidence that CTCF remains bound during meiosis ((Vara et al. 2019; Zuo et al. 2021a). A conditional CTCF KO mouse shows defects in spermatogenesis, first manifested in the pachytene stage (Hernández-Hernández et al. 2016). Therefore, CTCF might play a direct role in meiotic chromatin organization.

**Figure 4.**
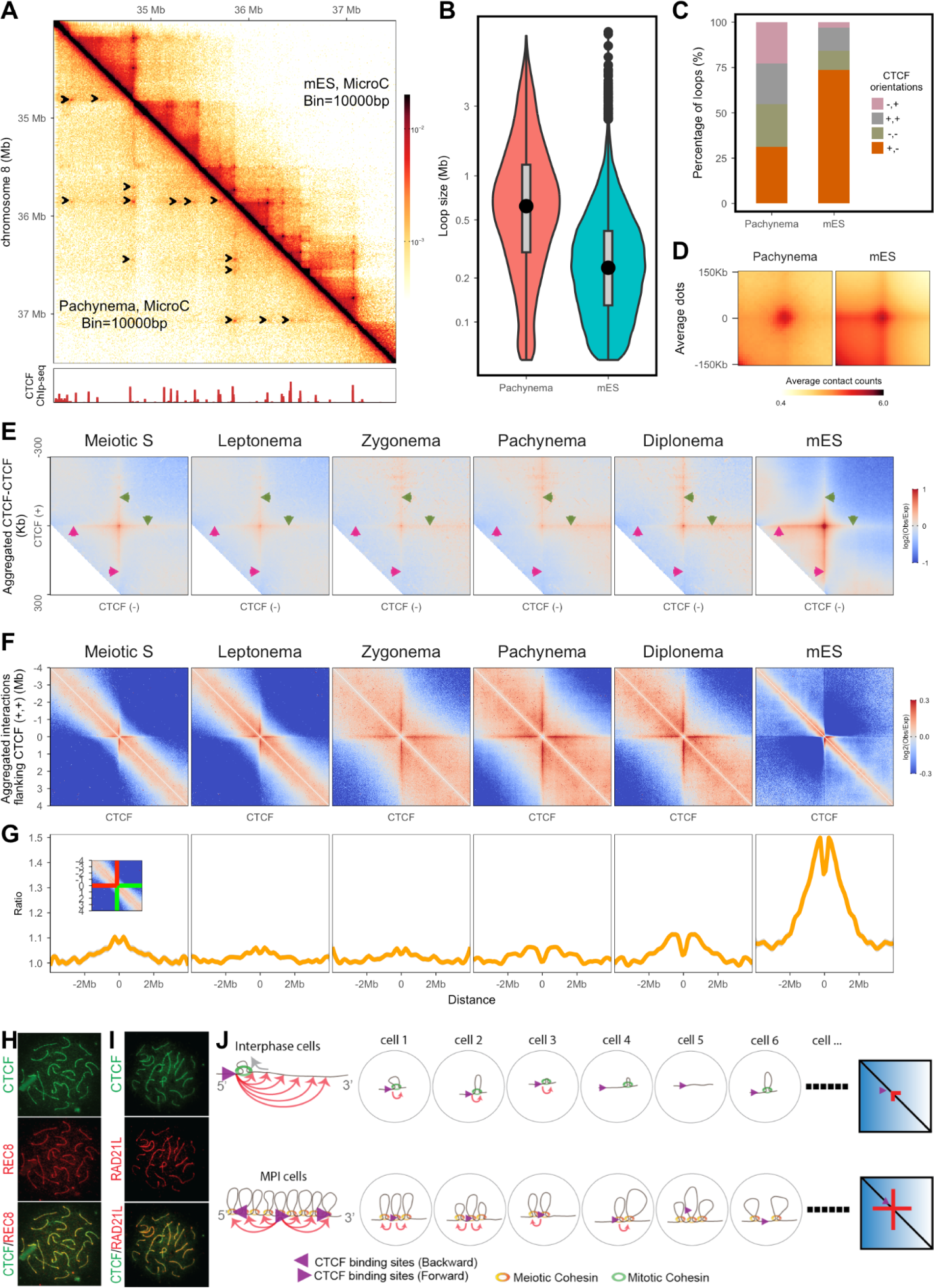
Meiotic specific interaction patterns shaped by CTCF and cohesin during meiotic prophase I. (A) Micro-C matrix of chromosome 8 at a resolution of 10 Kb (region: 34 Mb to 37.5 Mb). Upper half: mES, lower half: Pachynema. Both mES and pachytene Micro-C data were sequenced to a depth of ∼3 billion. Arrows indicate the dots located by Chromosight (Matthey-Doret et al. 2020). CTCF ChIP coverage is plotted at the bottom. (B) Median distance between anchors of identified dots in pachynema and mES cells. Dot calling was performed at both 5 Kb and 10 Kb resolution by chromosight (Matthey-Doret et al. 2020). (C) Percentage of dot anchors with different orientated CTCFs. Only both anchors containing one CTCF motif are considered for this analysis. (D) Aggregation of identified dots. The aggregation was performed at a resolution of 10 Kb, with a 100 Kb flanking region. (E) Aggregation of convergent CTCF interactions at a range of 100 Kb to 300 Kb. (F) Aggregation of interactions surrounding CTCF binding sites with unified motif orientation. The analysis was performed at a resolution of 10 Kb, with a 4 Mb flanking region. Only the interactions at loci with forward-oriented CTCF was shown. (G) Quantification of the orientation bias. The observed/expected values of each bin at the center of the stripes were extracted. The matrix is symmetric. Both green lines indicate the interactions between CTCF binding sites with the downstream loci; Both red lines indicate the interactions between CTCF binding sites with the upstream loci. The bias was calculated as the ratio between each value on the green line and the corresponding value on the red line. The profile was plotted symmetrically representing the symmetric tripes in the matrix. (H) A representative immunostaining image demonstrates that CTCF and REC8 are localized along the axial elements during the pachytene stage. (I) A representative immunostaining image demonstrates that CTCF and RAD21L are localized along the axial elements during the pachytene stage. (J) A schematic illustrates how multiple closely aligned loops and loop extrusion shape interaction patterns differently. The purple arrow represents genomic loci bound by forward-oriented CTCF. Orange curved lines with arrows point to possible contact loci. The possible looping status in each individual cell was drawn separately. As the loop bases are mostly random; and no difference would be anticipated on the two sides of a specific loop, CTCF bound loci might contact each nearby locus when resembling all interactions from multiple cells, resulting in an equal interaction enrichment in both directions as an average of a cell population. The depleted enrichment at short-distance indicates that all loops during MPI are above a certain size.

To better characterize these sites, we performed dot calling (see Methods) and identified 11,600 dots in pachynema, roughly half of the number identified from mES at the same sequencing depth (black arrowheads in Figure 4A, S12A). Approximately 80% of the dots feature at least one CTCF-defined anchor (Figure S12B). The median distance between two anchors of dots in pachynema is nearly 3 times bigger than that in mES cells (620 kb vs 235 kb) (Figure 4B). We found only 4% of pachytene dots to have identical anchor positions to those observed in mES cells. The dots in pachynema were “fuzzier” than the ones from mES after aggregating all pairwise signals (Figure 4D) indicating that the increased interactions extend to loci close to CTCF sites. Consistently with the disappearance of TADs, the increase in contacts in the region between two anchors is lost (Fig 4D). A major difference is that dots anchored between CTCF sites in any orientation were nearly equal in pachynema, whereas nearly 80% of dots in mES were anchored by convergent CTCF sites (Figure 4C).

We extended this analysis to all stages for which we have Micro-C data and called dots with the caveat that dot calling relies heavily on sequencing depth. The preference for dots being located at loci bound by CTCFs and the presence of more non-convergent dots were observed in all examined stages (Figure S12B-C). The progressive increase in distance between two anchors closely matches the trend in loop size in meiotic prophase I (Figure S5A-B, S12)

To further characterize the signal in each stage, we aggregated all interactions between CTCF binding sites (Figure S13A,S14) independently of dot calling. Given the progressive changes in loop size, pairwise CTCF were categorized based on both orientations and distances (Figure S14). Aggregation analysis confirmed the enrichment of interactions between CTCF binding sites at all orientations in all stages (Figure S13A, S14); and the intensities across these orientations were found to be similar (Figure S13A, S14).

In addition, we observed “stripes” emanating from the CTCF-CTCF interaction points; patterns are different depending on the distance between the CTCF sites (Figure S14; see Figure S15 for a schematic). Interestingly, a distinct dynamic was observed when aggregating the matrices around pairwise CTCF sites that are closer than 300 Kb: the stripe pattern in mES cells is compatible with increased interactions between either CTCF site and the DNA in the intervening region, proposed to be indicative of loop extrusion activities (Fudenberg et al. 2016; Vian et al. 2018; Mirny, Imakaev, and Abdennur 2019; Barrington et al. 2019; Park et al. 2023). In meiotic cells this pattern changes as cells progress into meiosis prophase I, the interactions between the CTCF sites and the DNA in between them is reduced while a strong interaction signal appears between the CTFC sites and the DNA flanking either site. This has been proposed to indicate an extension of pre-established loops (Haarhuis et al. 2017; van der Weide et al. 2021) in situations where cohesin is more stable.

Whether loop extrusion remains active during meiosis is under debate (Schalbetter et al. 2019; Castellano-Pozo et al. 2023; Patel et al. 2019; Wang et al. 2019), therefore we further explored the architectural “stripe” patterns in meiosis by aggregating contacts surrounding CTCF loci in the same orientation. A stripe pattern was observed in all cell types (Figure 4F). In mES cells the loci bound by forward-facing CTCF contact primarily with the downstream loci (Figure 4F,G). However such bias was largely lost throughout meiotic prophase I (Figure 4F,G). In pachynema, the interactions with loci upstream and downstream were nearly equal (Figure 4F,G) and they can extend to several megabases, significantly longer than the distance in mES cells (Figure 4F). Conversely, the enrichment at short distances was depleted (Figure S16) consistent with the existence of a preformed loop (see above). The orientation bias is a typical feature of architectural stripes indicative of loop extrusion (Fudenberg et al. 2017), the large reduction in this bias observed in meiotic cells suggested an alternative mechanism underlying the “stripe” pattern formation during meiosis. Recently, a “multi-loop” model was proposed from simulation, showing that the “stripe” pattern could also be the result of high density loading of cohesins (Chen et al. 2023). Closely positioned loops are a rare situation in interphase cells, nonetheless, it is how the chromatins are organized during meiosis (Figure 4J)

We found reproducible positioning of loops in our meiotic prophase data, suggesting that the interaction between CTCF and cohesins (Li et al. 2020) could establish an anchor point for the meiotic loop formation (Figure 4J). REC8 and RAD21L localize to the meiotic chromosome axis in meiotic prophase I (Ishiguro 2019). Colocalization of CTCF with the cohesin on the axes would support the notion that loci bound by CTCF are located at the base of the loop arrays at a high probability. A report analyzing a CTCF conditional deletion in meiosis showed a weak CTCF signal on the axes on pachytene spreads that disappeared in the KO cells (Hernández-Hernández et al. 2016). We prepared structurally preserved nuclei and spreads to revisit this question. We immunostained for CTCF using different antibodies and confirmed that CTFC is clearly bound to the axes as the synaptonemal complex assembles implicating CTCF in the organization of the meiotic loop array (Figure H-I).

Our data partially answers a long-standing puzzle: what are the sequence preferences located at the bases of meiotic loop arrays? While in yeast the positioning of loops depends on the presence of Rec8 and its DNA binding preferences, in mice, our data suggest that CTCF might play a role in stabilizing some of these cohesin complexes but it is not an absolute determinant of looping positions.

### The interactions between regulatory elements are well preserved during MPI

A challenge faced by meiotic chromatin is maintaining the balance between gene expression and the highly compacted genome. Taking advantage of our Micro-C data, we examined how the reorganization of MPI chromosomes influences interactions between regulatory elements.

We first built Micro-C matrices at 400 bp resolution, which enabled us to observe the self-interaction domains at the scale of single gene units (Figure 5A, Figure S17). These domains encompass either single or multiple genes, in addition to intergenic regions. These observations echo findings from mES cells (Figure 5A, Figure S17; (Hsieh et al. 2020)). Additionally, our Micro-C maps hinted at the presence of stripes emanating from transcription start sites (TSSs) and enriched interactions between TSSs throughout meiotic prophase I. However, the signals from the matrix snapshot are faint, preventing us from drawing any definitive conclusions.

**Figure 5.**
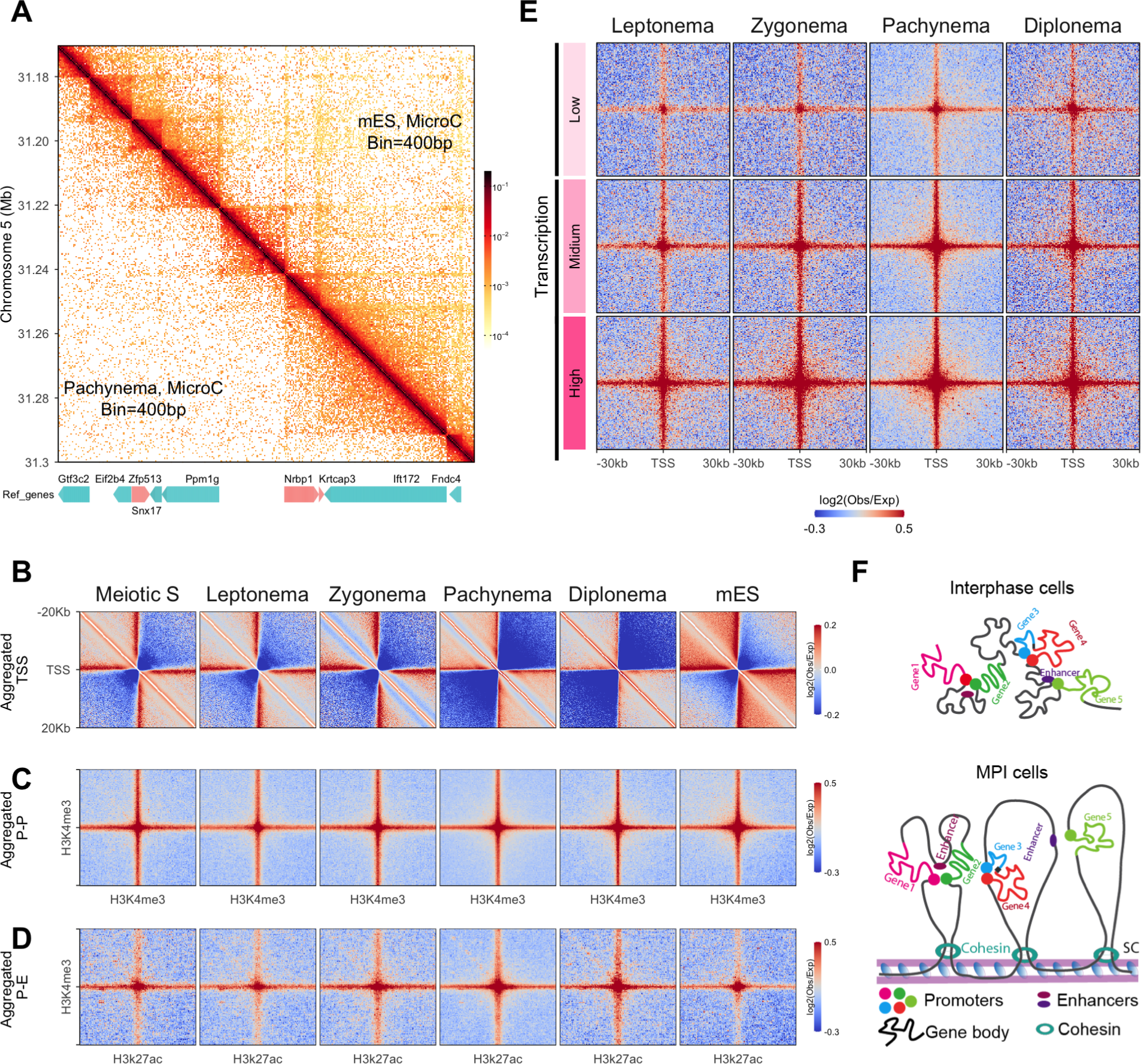
Micro-C revealed the preservation of fine-scale chromatin structure during MPI. (A) A snapshot of a Micro-C matrix of chromosome 5 at a resolution of 400 bp (region: 31 Mb to 31.4 Mb). Upper half: mES, lower half: Pachynema. Genes in the same region were annotated at the bottom. The arrows on the bars indicate the orientation of the genes. (B) Aggregated interactions centered around TSSs. TSSs were derived from USCS annotated genes. The aggregated interactions were calculated at a resolution of 200 bp with a flanking region of 20 Kb. (C). Aggregated promoter-promoter interactions. The promoters were derived from H3K4me3 Chip-seq data (Lam et al. 2019). The H3K4me3 modification sites that overlapped with hotspots were excluded from calculations. Resolution was 400 bp with a flanking region of 30 Kb. (D) Aggregated promoter-enhancer interactions. The enhancers were derived from H3K27ac Chip-seq data (Lam et al. 2019);(Maezawa et al. 2020). Common peaks from both studies were selected and only the sites that did not overlap with H3K4me3 peaks and hotspots were used for the calculation. Resolution was 400 bp with a flanking region of 30 Kb. (E) The correlation between gene expression and P-P interactions. The genomic loci with H3K4me3 modification were divided into three groups based on their expression deduced from RNA seq data (Gaysinskaya et al. 2018). High: Represents the top third in terms of expression levels across all genes; Mid: Corresponds to the middle third in expression levels among all genes; Low: Encompasses the remaining genes, covering the lowest third in expression levels. The aggregated interactions were calculated at a resolution of 400 bp with a flanking region of 30 Kb. (F) Schematic of fine-scale structures from MPI. Top: Somatic interphase structures were adapted from (Hsieh et al. 2020). Bottom: Potential fine-scale structures within the MPI loop array.

To validate these observations, we aggregated the interactions flanking TSSs at a resolution of 200 bp using our Micro-C data. This analysis revealed stripe signals persisted throughout all examined stages, with comparable intensity to that of mES cells (Figure 5B). In line with a previous study with mES, the formation of these stripes was independent of CTCFs (Figure S18B; Hsieh et al. 2020). Interestingly, we found that the insulation between genes was increased at pachynema and diplonema (Figure 5B, Figure S18A,B). Given the pachynema we collected is relatively late, this phenomenon could potentially be the result of a significant accumulation of “sticky” RNA Pol II on genes at these stages (Alexander et al. 2023).

To further examine the interactions between regulatory elements, we aggregated the pairwise promoter-promoter (P-P; H3K4me3-H3K4me3) and promoter-enhancer (P-E; H3K4me3-H3K27ac) interactions. We observed clear P-P and P-E interactions at all stages, with similar intensity as mES cells (Figure 5C,D). These interactions were not observed from Hi-C data and are independent of CTCFs (Figure S19A,B; Figure S20). Furthermore, in line with observations from interphase cells (Friman et al. 2023), we found that the P-P and P-E interactions could occur between two loci that are extremely distant from one another (Figure S21). Unlike interphase cells, we observed a marked decrease in ultra-long-distance interactions (beyond 30 Mb) from zygonema to diplonema (Figure S21A,B). Even though the communication of regulatory elements on different chromosomes has been observed in interphase cells, such interactions were barely visible after zygonema during MPI (Figure S22). To further understand the regulatory role of these interactions, we categorized the promoters based on expression level (Gaysinskaya et al. 2018) and aggregated their interactions. This analysis demonstrated that the P-P interactions were associated with transcriptional levels, indicating these interactions may play a regulatory role (Figure 5E).

Our data implies that despite the MPI chromatins being highly compacted by the formation of loop arrays, the fundamental interactions between regulatory elements were well preserved. However, the interactions are regulated by the meiotic-specific genomic configuration (Figure 5F).

## Discussion

For the proper chromosomal separation, both meiotic and mitotic chromosomes undergo compression through loop array formation (Kleckner, Zickler, and Witz 2013; Gibcus et al. 2018; Zickler and Kleckner 2023). While mitotic chromosomes lose most of the interphase features (Gibcus et al. 2018), certain characteristics, although altered, are preserved on meiotic prophase I chromosomes. This might be associated with the additional challenges faced by meiotic chromosomes, such as the execution of the meiotic recombination program and active transcription maintenance. Here, by examining the genomic interactions at the highest temporal and spatial resolution to date, we gained insights into the principles of chromosomal organization during mammalian meiosis.

We show that long-range whole-genome compartmentalization is lost consistent with the formation of a loop array. Nonetheless, subcompartments are preserved locally. Previous studies identified refined local compartments, but they were not observed in the corresponding regions of interphase cells (Patel et al. 2019; Alavattam et al. 2019; Wang et al. 2019, (He et al. 2023) leading to the conclusion that they were unique features of meiotic chromosomes. By examining deep-sequenced Hi-C matrices from stages preceding meiosis, we found that the local “checker-board”-like pattern was also present in germline interphase cells, although with various interaction strengths. Importantly, a recent study revealed that the median size of compartmental interval in somatic interphase cells is much finer than it was assumed, with a median size of 12.5 Kb (Harris et al. 2023). In addition, this study showed that all active transcriptional regulatory elements interact within the A compartment (Harris et al. 2023). A striking feature of the Hi-C maps as cells enter meiosis is the disappearance of TADs. We propose that the local compartments are more visible in meiosis due to changes on cohesin activities like loop extrusion that would counteract these interactions. Overall, the similarity of compartment intervals between germline interphase and prophase cells point to a common underlying principle governing chromosome organization in these two cell types.

We isolated a rare cell population that had left mitotic cycles yet not entered meiosis. These transition stages enabled us to uncover an unexpected shift in chromatin folding. We found that chromosomes at that stage are mostly devoid of loops. This initial chromatin reorganization coincides with a decrease in protein levels of the mitotic cohesin and a concomitant increase in the meiotic cohesin complexes. As cells progress into meiosis, an increased number of extended loops are formed, which coincide with the increase in REC8. Interestingly, the cohesin exchange is accompanied by the establishment of a folding intermediate which was only observed when somatic cells exit mitosis (Abramo et al., n.d.). Whether this intermediate folding state is conserved when switching between SMC complexes (cohesin to condensin or mitotic to meiotic cohesin) needs to be further examined.

Interestingly, our analyses from transition stages I and II revealed similarities to the dynamics of loop formation in WAPL-deficient cells. Losing the cohesin unloader WAPL results in stronger corner peaks at TAD boundaries (Hi-C loops) and a relative increase in inter-TAD interactions (Haarhuis et al. 2017). It was proposed that WAPL-mediated removal of cohesin, as the loops are forming between two CTCF sites, contribute to the increase of intra-TAD interactions and the consequent high insulation score. The average loop size was also extended resembling our observations from the transition stages. This suggests that these features are common to situations where the cohesin complexes are more stable.

There is no evidence to date for DNA sequence specificity in loop positioning in mammals. Previous Hi-C experiments failed to detect any distinct looping positions in mammals (Grey and de Massy 2021). We combined Micro-C experiments with high sequencing depth to examine this question. We found a clear pattern in meiotic prophase I consistent with well positioned loops. These “dots” are reminiscent of the punctate grid-like interactions described in yeast that are mediated by the meiotic cohesin subunit, Rec8 (Schalbetter et al. 2019). We found here that they coincide with CTCF sites. Importantly, distinct from mitotic cells, they do not show increased interactions in the intervening regions, suggesting that they are not the typical TAD corner signal. Furthermore, convergent CTCF motifs are not predominantly found at these sites. We show that CTCF colocalizes with REC8 at the chromosome axes supporting the notion that the dots we identified are at the base of the bonafide meiotic loops. We cannot distinguish whether all loops have CTCF at their bases but there is a clear preference for this configuration. Indeed we found that up to 20% of “dots” in pachymena do not have a CTCF binding site at their anchors. The densely loaded cohesins may act as anchors independently of CTCF in some cases. Furthermore, we observed that unlike in interphase cells, dots look “fuzzier” meaning that there are interactions beyond the precise CTCF binding site, maybe reflecting the more dynamic and/or flexible nature of these interactions that don’t require convergent CTCF sites. Having a flexible but reproducible looping point might be an important feature that allows both homologs to share a loop domain that in turn would facilitate homology search and double-strand break repair.

In interphase cells, the association and dissociation of cohesins is transient, and emerging live-cell microscopy data found that loops are both rare and short-lived (Gabriele et al. 2022; Mach et al. 2022). During meiotic prophase I, the loop array remains in place for days but how they form and “grow” is unclear. Our data cannot formally distinguish between active loop extrusion or multiple loops forming statically and then fusing. We favor the hypothesis of loops forming statically rather than dynamically due to the presence of corner dots without TADs. Therefore, the meiotic loop array might be dependent on the stability of the cohesin complexes rather than its loop extrusion activities. The cohesin regulators, WAPL and NIPBL are expressed through mammalian meiosis, and perturbing this regulation would inform how the stability of cohesins affects the meiotic structure and function.

Finally, our data revealed a unique global chromatin organization that results from balancing between the dense linear loop array configuration and maintaining active transcription. The interactions between regulatory elements, key factors in regulating gene transcription, are well preserved. This is in line with the observation that P-P/P-E interactions are resilient to acute loss of architectural factors, such as CTCF, cohesin, WAPL, YY1 and RNA polymerase II (Hsieh et al. 2022; Goel, Huseyin, and Hansen 2023). Two models were proposed regarding the establishment and maintenance of P-P/P-E interactions. In the time-buffering model, chromatin architectural factors are required for establishing these interactions, rather than maintaining them (Hsieh et al. 2022). Whether the transcriptional program is pre-set before entering meiosis remains to be answered. In a more recent study, the deepest interaction map was generated by region capture Micro-C (Goel, Huseyin, and Hansen 2023). Based on the observed highly nested fine-scale interactions, these contacts between regulatory elements were proposed to be driven by microphase separation, resembling A/B compartmentalization (Goel, Huseyin, and Hansen 2023). The two models are not mutually excluded. The P-P/P-E interactions during MPI potentially follow the same rule. Importantly, it is not clear what structural features are relevant for gene regulation. Here we show that in the absence of TADs and within a highly structured loop array, the transcriptional program is not affected.

The development of new technologies has allowed us to interrogate the establishment of meiotic chromatin in unprecedented detail. Although many questions remain, the framework established in this work will pave the way toward a better understanding of the interplay between structure and function.

## Methods

### Mouse strain

C57BL/6J (Stock no. 000664) and CAST/EiJ (Stock no. 000928) were purchased from The Jackson Laboratory. The F1 hybrids were in-house bred with male CAST/EiJ with female C57BL/6J.

### Antibodies used

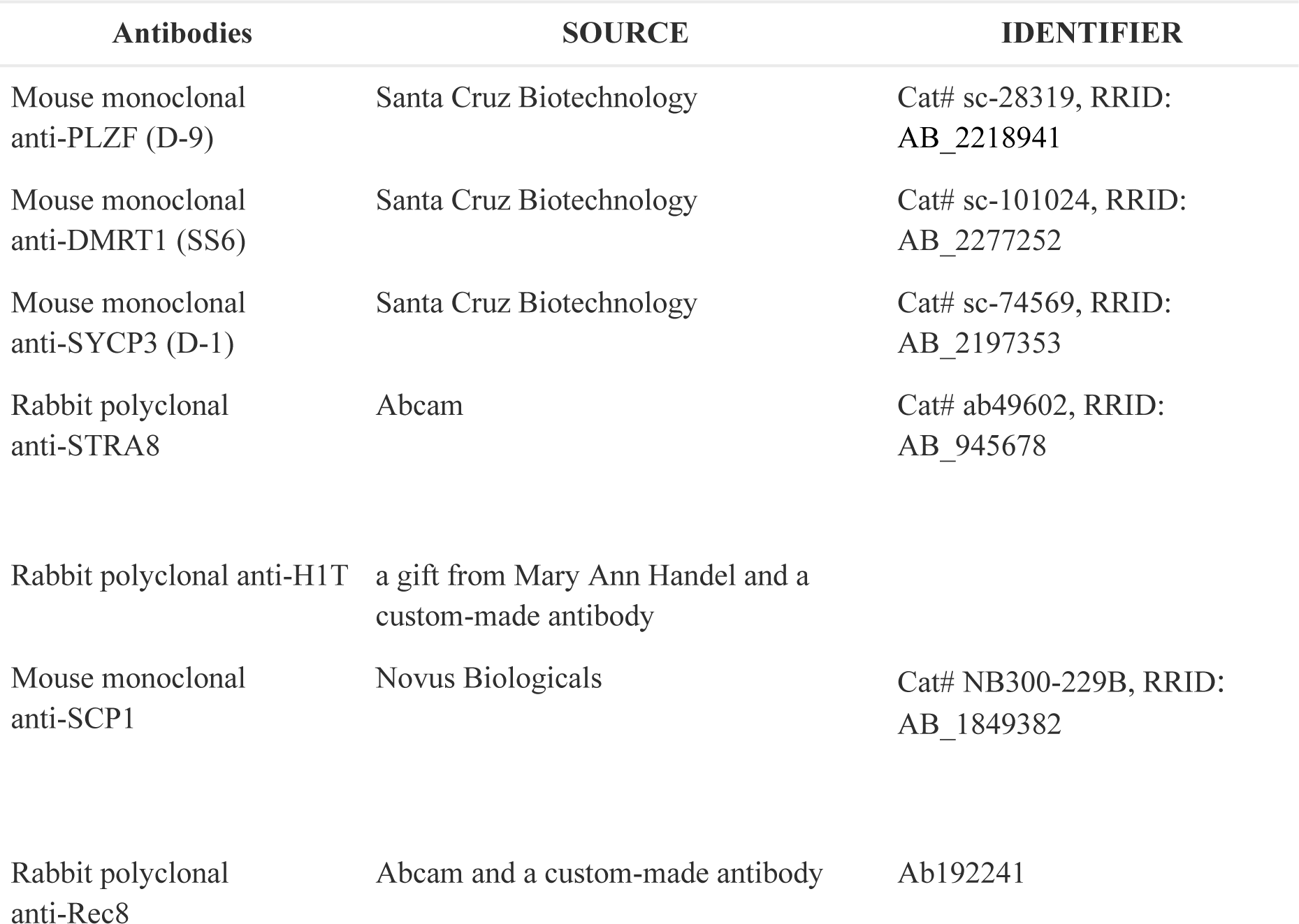
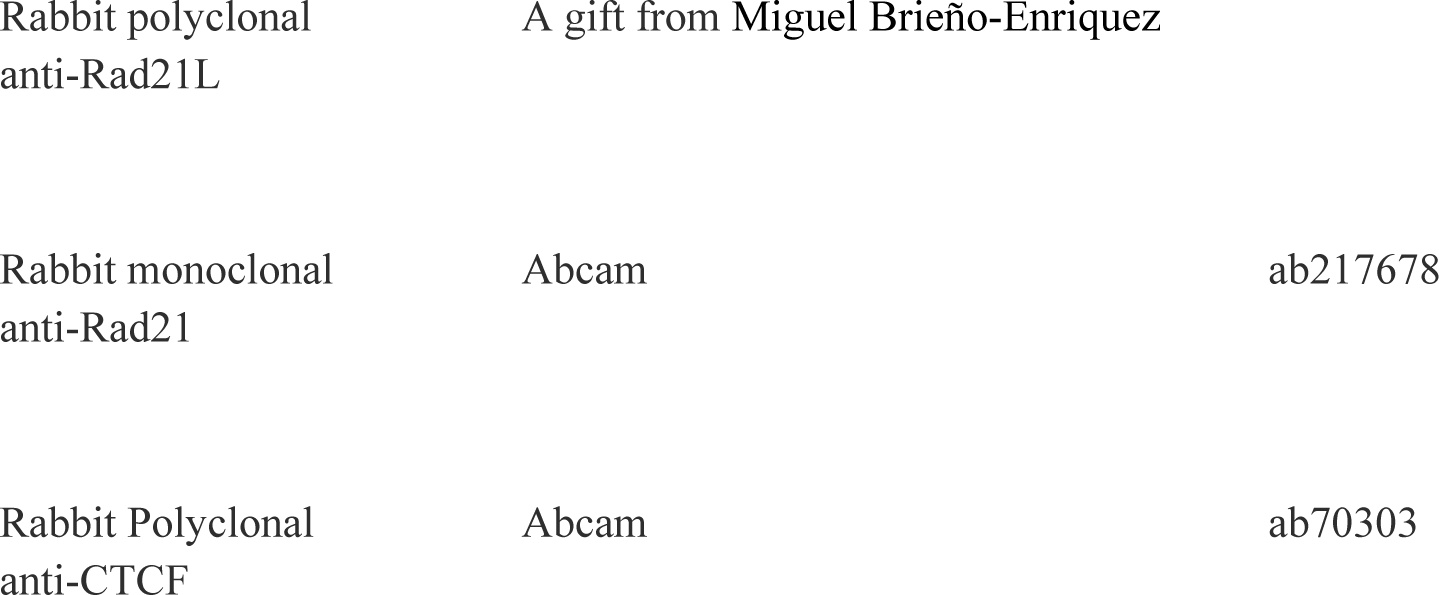

| Antibodies | SOURCE | IDENTIFIER |
| --- | --- | --- |
| Mouse monoclonal anti-PLZF (D-9) | Santa Cruz Biotechnology | Cat# sc-28319, RRID: AB_2218941 |
| Mouse monoclonal anti-DMRT1 (SS6) | Santa Cruz Biotechnology | Cat# sc-101024, RRID: AB_2277252 |
| Mouse monoclonal anti-SYCP3 (D-1) | Santa Cruz Biotechnology | Cat# sc-74569, RRID: AB_2197353 |
| Rabbit polyclonal anti-STRA8 | Abcam | Cat# ab49602, RRID: AB_945678 |
| Rabbit polyclonal anti-H1T | a gift from Mary Ann Handel and a custom-made antibody |  |
| Mouse monoclonal anti-SCP1 | Novus Biologicals | Cat# NB300-229B, RRID: AB_1849382 |
| Rabbit polyclonal anti-Rec8 | Abcam and a custom-made antibody | Ab192241 |
| Rabbit polyclonal<br>anti-Rad21L | A gift from Miguel Brieño-Enriquez |  |
| Rabbit monoclonal<br>anti-Rad21 | Abcam | ab217678 |
| Rabbit Polyclonal<br>anti-CTCF | Abcam | ab70303 |

### Nuclei preparation and isolation by FACS

The nuclei were prepared and sorted as described before (Lam et al. 2019). Briefly, the dissected testes were first fixed with 1% formaldehyde for 10 minutes for Hi-C and Chip-seq (or 1% formaldehyde for 10 minutes + disuccinimidyl glutarate for 45 minutes). The fixatives were quenched by 0.125 M glycine and homogenized. The cell suspension was filtered by 70 μm cell strainer, washed with 1X PBS (phosphate buffered saline) and then resuspended in nucleus extraction buffer (15 mM Tris-HCl pH 7.4, 0.34 M sucrose, 15 mM NaCl, 60 mM KCl, 0.2 mM EDTA (Ethylenediaminetetraacetic acid), 0.2 mM EGTA (ethylene glycol tetraacetic acid). Nuclei were extracted using a Dounce homogenizer, with 30 strokes with the tight pestle, and repeated once. Nuclei were filtered through a 40-μm cell strainer and resuspended in a chilled PBTB buffer (1× PBS with 0.1% Triton X-100, 5% bovine serum albumin, and protease inhibitor).

The nuclei were incubated with primary antibodies at room temperature for 1 hour or 4 °C overnight. They were then washed with PBTB and labeled with secondary antibodies at room temperature for 30 minutes. After washing twice with PBTB, nuclei were filtered with a 40-μm cell strainer. The nuclei were stained with DAPI before sorting. Singlets were gated by FSC and FAC. 2C, 2C-4C, and 4C populations were first separated by DAPI signal. Then targeted populations were isolated by the combination of different markers. The sorted nuclei were collected in PBTB. The collected nuclei were either used for downstream experiments or stored at -80 °C.

### Micro-C

Micro-C was performed as described(Hsieh et al. 2020) with minor modifications. 0.5 million sorted nuclei were resuspended with 100 μL MBuffer#1 (50mM NaCl, 10mM Tris-HCl pH 7.5, 5mM MgCl2, 1mM CaCl2, 0.2% NP-40, 1x protease inhibitor cocktail (Roche, 04693132001) and incubated on ice for 20 min. The nuclei were collected by centrifugation (900g, 5 min) and resuspended with 100 μL MBuffer#1. Added 2.5 U MNase (Worthington Biochem #LS004798) to the solution and incubated at 37°C for 10 min. Add 2 μL 200 mM EGTA and incubate at 65°C for 10 min to stop the reaction. Wash/rinse the nuclei with 1xNEBbuffer2.1(NEB, #B7202S) for three times, and resuspended the nuclei with 45 μL 1xNEBbuffer2.1. Dephosphorylation of DNA ends was performed by addition of 5 μl rSAP (NEB, #M0203) and incubation at 37°C for 45 min. rSAP was inactivated by incubating at 65°C for 5 min. Add pre-mix chewing solution (5 μl 10xNEBbuffer2.1, 3 μl 100 mM DTT (ThermoFisher, A39255), 2 μl 100 mM ATP (ThermoFisher, R0441), 30 μl H2O), 8 μl 5 U/μL Klenow Fragment (NEB, M0210S) and 2 μl T4 PNK (NEB, #M0201L) to the dephosphorylated nuclei sample and incubate at 37°C for 15 min. The end repair and Biotin labeling was performed by addition of (25 μl 0.4 mM Biotin-dATP, 25 μl 0.4 mM Biotin-dCTP, 1+1 μl 10 mM dTTP + dGTP, 10 μl 10XT4 DNA Ligase Buffer and 1 μl 10 mg/mL BSA) and incubated at 25°C for 45 min. The reaction was stopped by addition of 12 μl 0.5 M EDTA and incubation at 65°C for 20 min. The nuclei were collected by centrifugation (900g, 5 min). Wash/rinse the nuclei with 1xNEB ligation buffer with 0.1% Triton X-100. The nuclei were resuspended with ligation-mix (100 μl 10X T4 DNA Ligase Buffer, 5 μl 20 mg/mL BSA, 1 μl 20000 U/mL T4 DNA Ligase, 100 μl 0.1% Triton X-100, 884 μl H2O) and incubated at room temperature for 3 hours. To remove the unligated biotin, the ligated nuclei were pelleted and resuspended with 200 μl 1xNEB buffer1 with 200 U Exonuclease III and incubated at 37°C for 5 min. Reverse crosslinking was performed by addition of 25 μl proteinase K and 25 μl 10% SDS, then incubation at 65°C overnight. The DNA was purified by phenol/chloroform and then fragments above 200 bp were selected by AMPure XP (BECKMAN COULTR, A63881). The end repair and adaptor ligation was performed by KAPA HyperPrep Kit (Roche, 07962347001). Adapter ligated DNA was purified by AMPure XP and eluted with 100 ul EB buffer (Qiagen, 19086). 10 μl Dynabeads™ M-280 Streptavidin (ThermoFisher, 11205D) for each sample was washed three time with 2xB&W Buffer (10 mM Tris-HCl (pH 7.5), 1 mM EDTA, 2 M NaCl) and resuspended with 100 μl 2xB&W Buffer. The beads were added to adaptor ligated DNA and incubated for 15 minutes. Biotin labeled DNA was attached to the beads. The beads were then washed by 1xTB&W (5 mM Tris-HCl (pH 7.5), 0.5 mM EDTA, 1 M NaCl, 0.05% tween 20) for three times, and washed by 10 mM Tris-Cl pH 8.0 once. The beads were resuspended with a 20 μl EB buffer. PCR was performed by KAPA HyperPrep Kit. A size selection was performed by loading the amplicon to 1.5% agarose gel and extracting the band of about 400 bp. The cut band from the gel was then purified by DNA Recovery Kit (Zymo Research, #D4002).

### Hi-C

Hi-C experiments were performed as described (Belaghzal, Dekker, and Gibcus 2017) with some modifications. The sorted nuclei were resuspended with a digestion buffer (44 ul of 10x NEB DpnII buffer, 38 μl of 1% SDS, and 300 ul H2O) and incubated at 65 °C for 10 minutes. Quench the SDS with by addition of 44 ul 10% Triton X-100 and incubate at 37 °C for 1 hour. The chromatin was digested by addition of 400 U DpnII and incubated at 37 °C overnight (12 hours to 16 hours). 10 ul of the mixture was used for quality control of the digestion. The digested nuclei were washed with 1x buffer2 once and resuspended with premixed biotin fill-in buffer (1.5 ul 10 mM dCTP, 1.5 ul 10 mM dGTP, 1.5 ul 10 mM dTTP, 37.5 ul 0.4 mM biotin-14-dATP, 40 ul NEB buffer2, 10 ul 5U/ul DNA polymerase I Klenow) and incubate at 37 °C for 1 hour. The biotin labeled nuclei were washed once with 1x NEB ligation buffer and resuspended with ligation mix (100 ul 10x NEB ligation buffer, 100 ul 10% Triton X-100, 10 ul 10 mg/ml BSA, 2 ul 2000 U/ul T4 DNA ligase) and incubated at room temperature for 4 hours. The ligated nuclei were pelleted and resuspended with a reverse crosslink buffer (150 ul H2O, 5 ul 5M NaCl, 5 ul 10 mg/ml Rnase). The reverse cross-linking was performed by addition of 20 ul 10% SDS and 20 ul 20 mg/ml proteinase K and incubated at 65 °C overnight. The DNA was purified by phenol:chloroform then chloroform. The DNA pellet was resuspended with 50 ul EB buffer (Qiagen).

### ChIP-seq

Resuspended the nuclei with lysed with Lysis buffer 1 (0.25% Triton X-100, 10 mM EDTA, 0.5 mM EGTA, 10 mM Tris-HCl pH 8) and incubated at room temperature for 10 min. The nuclei were collected by centrifugation (900g, 5 min) and resuspended with Lysis buffer 2 (200 mM NaCl, 1 mM EDTA, 0.5 mM EGTA, 10 mM Tris-HCl pH 8). Then collect the nuclei by centrifugation (900g, 5 min) and resuspended with RIPA buffer (10 mM Tris-HCl pH 8, 1 mM EDTA, 0.5 mM EGTA, 1% Triton X-100, 0.1% sodium deoxycholate, 0.1% SDS (sodium dodecyl sulfate), 140 mM NaCl plus protease inhibitor). Chromatin was sheared into ∼50–300 bp fragments by sonication using Bioruptor (Diagenode).

The immunoprecipitation was performed by addition of 1-2ug antibodies and incubated at 4 °C overnight. To pull down the immuno-complexes, add 50 μl Dynabeads Protein G (30 mg/ml, Novex) and incubate at 4 °C for 2 hours. Wash the bead once with low salt buffer (0.1% SDS, 1% Triton-X-100, 2 mM EDTA, 20 mM Tris-HCl pH 8, 150 mM NaCl), once with high salt buffer (0.1% SDS, 1% Triton-X-100, 2 mM EDTA, 20 mM Tris-HCl pH 8, 500 mM NaCl), twice with LiCl buffer (0.25 M LiCl, 1% IGEPAL-CA630, 1% sodium deoxycholate, 1 mM EDTA, 10 mM Tris-HCl pH 8) and twice with TE buffer. The ChIPed DNA was purified following the protocol of IPure kit v2 (diagenode). Libraries are prepared by KAPA HyperPrep Kit (Roche, 07962347001). DNA sequencing was performed on the Illumina HiSeq 2500 or HiSeq X.

### Hi-C/Micro-C data processing

Both Hi-C and Micro-C data are processed following the 4DN Hi-C data processing pipeline. Briefly, the pair-end sequenced reads are mapped to mm10 using BWA (version 0.7). The pairs were then filtered by Pairtools (Open2C et al. 2023) to generate valid pairs. Matrices at different resolutions were built and balanced by the Cooler package (Abdennur and Mirny 2020).

### ChIP-seq analysis

The sequenced tags were mapped to the mouse mm10 reference genome with BWA-MEM(H. Li 2013). Peaks for CTCF and H3K27ac ChIP-seq were called using MACS2(version 2.2.7.1(Yong Zhang et al. 2008)) with default parameters. Peaks for H3k4me3 ChIP-seq performed with substages of MPI were called in our previous study(Lam et al. 2019) and merged in this study.

Two H3K27ac ChIP-seq data were used in this study. The first one is from our previous study, which was performed on SCP3^postive^,H1t^negative^ sorted nuclei. The second one is from Dr. Satoshi H. Namekawa’ group (Maezawa et al. 2020), which is performed on spermatocytes. The common peaks from both studies, excluding the peaks that overlap with H3K4me3 peaks, were used as enhancers.

CTCF motifs were called using FIMO from MEME suite (version 5.4.1)(Bailey et al. 2015) with JASPAR pwm MA0139. All parameters were set as default.

### Contact probability decaying curve and derivatives

The contact probability as a function of genomic distance *P*(s) curves on 1 Kb binned data were computed using cooltools.expected_cis from Cooltools (version 0.5.4)(Open2C et al. 2022). The curves were smoothed at log-scale on both y-axis and x-axis. Given the special configuration of chrX and chrY during MPI, only auto chromosomes were considered in this calculation. The derivatives were calculated at log-log scale from the smoothed *P*(s) curves.

### Insulation and TAD analysis

We calculated insulation score (Crane et al. 2015) at 10 Kb resolution with sliding window 100 Kb by Cooltools (Open2C et al. 2022). To call the boundaries, a threshold was set by threshold="Li", and regions with stronger insulation than the threshold were identified as boundaries. TAD domains were called as regions between two adjacent boundaries.

To measure the average insulation between TADs, we first identified conserved TAD boundaries by overlapping the TAD boundaries (± 10 Kb expansion) from unDiff.SGA and mES (Hi-C data from (Yan et al. 2018)). The average insulation score of the conserved TAD boundaries and the flanking regions (± 150 Kb from TAD boundaries) were calculated. To enable a standardized comparison of insulation scores across the analyzed stages, the values were normalized by assigning “zero” to the TAD boundary.

To reveal the dynamics of TADs, we identified the conserved TAD domains by selecting the domains called from unDiff. SGA and defined by conserved boundaries. The aggregated computation is performed by Cooltools(Open2C et al. 2022)

### Hi-C loop calling

The Hi-C loops are called by Chromosight (Matthey-Doret et al. 2020) at resolutions of 5 Kb and 10 Kb with the following parameters: --min-dist 50000 --max-dist 10000000 --min-separation 10000 --pearson=0.4.

### Compartment analysis

PCA based method and Calder (Y. Liu et al. 2021) were both used to identify compartment domains. Eigenvector decomposition was performed at a resolution of 100 Kb using eign-cis from Cooltools (version 0.5.4) (Open2C et al. 2022). GC content was used to orient the first eigenvector. Compartments calculated by Calder was performed at 50 Kb resolution with default parameters. The compartment calculations were carried out by the Calder algorithm at a 50 Kb resolution using default settings

### Aggregated analysis

All aggregated interactions in this study were calculated using Coolpuppy (version 1.0.0) (Flyamer, Illingworth, and Bickmore 2020) and plotted in R.

To aggregate the interactions between CTCFs, we selected the CTCF binding sites that are both at the top quartile of peak strength and the quartile with the lowest p-values in motif calling. The motifs with different orientations were aggregated separately. If not specified otherwise, pairwise CTCF binding sites positioned within a range of 100 Kb to 2 Mb were used for calculation. To measure the enrichment at different stages, we calculated the average of the three central pixels.

The aggregation of P-P and P-E interactions were conducted at sites positioned within a range of 5 Kb to 5 Mb, if not specified.

## ACKNOWLEDGMENTS

We thank Galina Petukhova and Michael Lichten for critiques; the Flow Cytometry Core of NHLBI; the NIDDK Laboratory of Animal Sciences Section; and the NIDDK genomics core facility. This work utilized the computational resources of the NIH HPC Biowulf cluster (https://hpc.nih.gov). This work was funded by the NIDDK intramural program (to R.D.C.O.).

