## supplementary materials for "High resolution maps of chromatin reorganization through mouse meiosis reveal novel features of the 3D meiotic structure"

### Supplementary figures

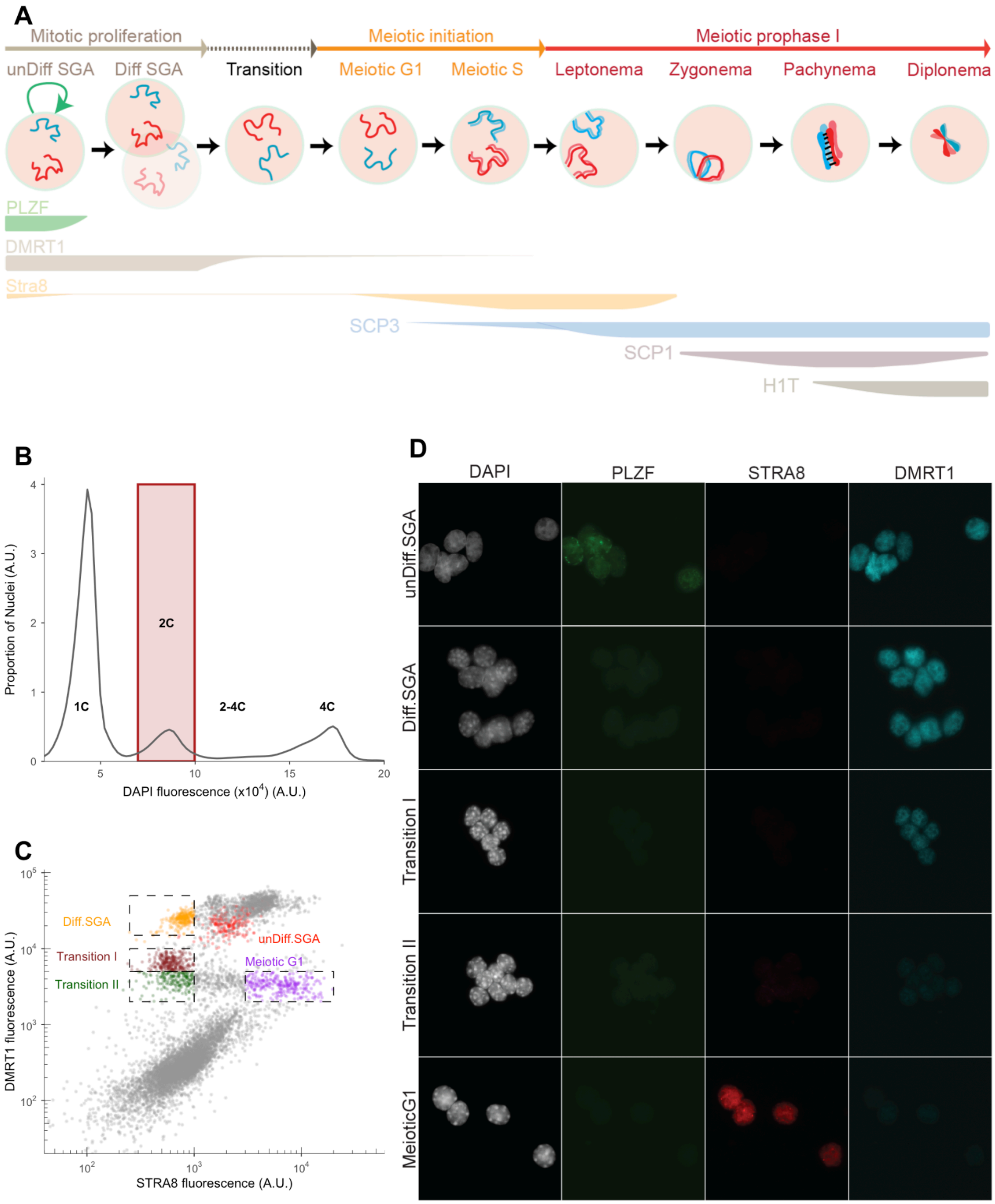

**Figure S1. Isolation of stage specific nuclei through spermatogenesis.**

(A) A schematic of the strategy for isolating nuclei from each stage. Intra-nuclear markers used for sorting were labeled with different colors. The relative expression level of each protein is represented as the thickness of their corresponding lines. Nuclei from unDiff.SGA to the Meiotic G1 stage were isolated with a combination of DAPI, PLZF, DMRT1 and STRA8. Nuclei from the Meiotic S stage to Leptonema were isolated with a combination of DAPI, SCP3, DMRT1 and STRA8. Nuclei from the leptonema stage to diplonema were isolated with a combination of DAPI, SCP3, H1T1 and SCP1.

(B-C) Strategies for isolating nuclei from unDiff.SGA to the Meiotic G1 stage.

(B) Nuclei from unDiff.SGA to Meiotic G1 stage were isolated from the 2C population.

(C) A snapshot from flow cytometry illustrating the gating for each population. Undifferentiated spermatogonia (unDiff SGA) were identified as 2C nuclei expressing the PLZF (promyelocytic leukemia zinc finger) protein (Buaas et al. 2004; Costoya et al. 2004). Other “pre-meiotic” populations were identified using a combination of DMRT1, which suppresses meiotic entry and is expressed throughout spermatogonial development (Matson et al. 2010), and STRA8, which can trigger meiotic entry but is expressed both in spermatogonia (Zhou et al. 2008; Endo et al. 2015) and in cells that enter MPI (Endo et al. 2015). We identified three populations of nuclei along the continuum of expression of these proteins to represent a putative commitment trajectory of mitotic germ cells as they entered meiosis. The differentiating spermatogonia (Diff.SGA) were collected as nuclei that expressed high levels of DMRT1 and no STRA8. The mitotic-to-meiotic transition nuclei were identified as a population with reduced DMRT1 and slightly increased STRA8. To further examine the transition between mitotic and meiotic stages, the transition population of nuclei was separated into Transition I and Transition II stages based on the DMRT1 signal. Finally, the nuclei with increased STRA8 and reduced expression of DMRT1 were collected from the 2C population as Meiotic G1. PLZF+ = unDiff.SGA; P2= Diff.SGA; P3= Transition I; P4= Transition II; P5= MeioticG1.

(D) Representative images of isolated nuclei from unDiff.SGA to the Meiotic G1 stage.

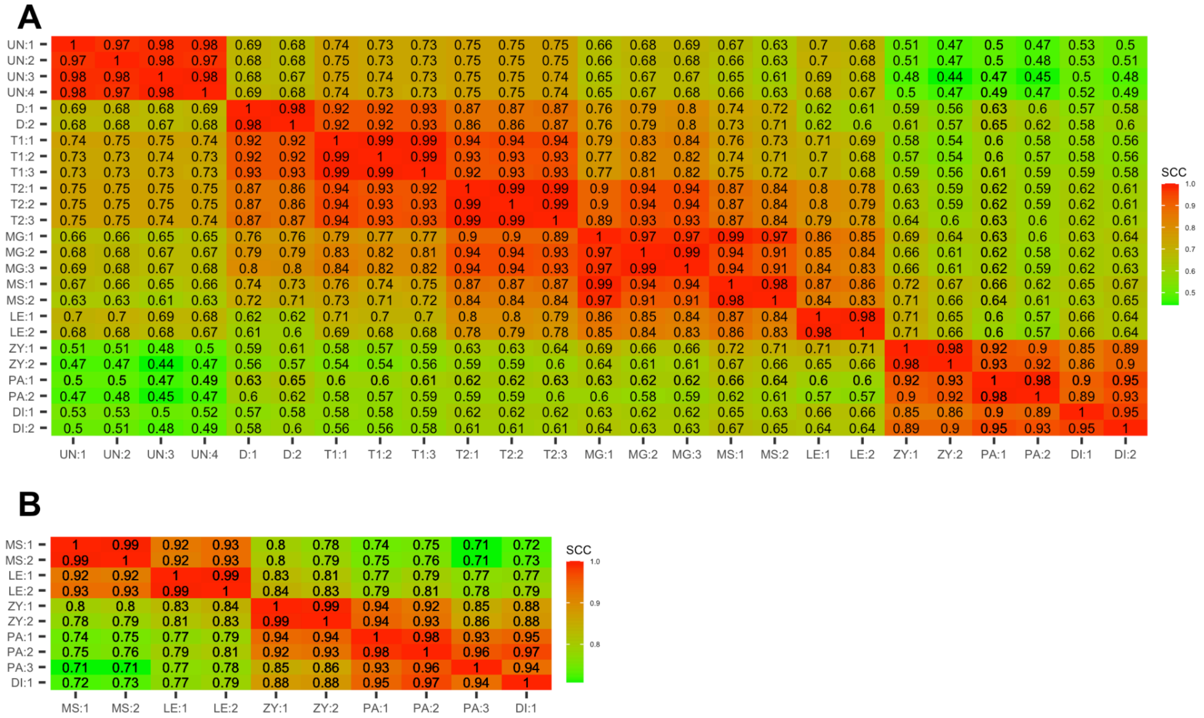


**Figure S2. Reproducibility of the biological replicates.**

(A) Reproducibility of Hi-C data.

(B) Reproducibility of Micro-C data.

The reproducibility was assessed by HiCRep (Yang et al. 2017; Lin, Sanders, and Noble 2021). The stratum-adjusted correlation coefficient (SCC) was calculated with a smooth parameter h=11, bin size=25,000 and maximal genomic distance to include in the calculation dBPMax = 500000.

UN=unDiff.SGA; D=Diff.SGA; T1=Transition I; T2=Transition II; MG=MeioticG1; MS=Meiotic S; LE=Leptonema; ZY=Zygonema; PA=Pachynema; DI=Diplonema.

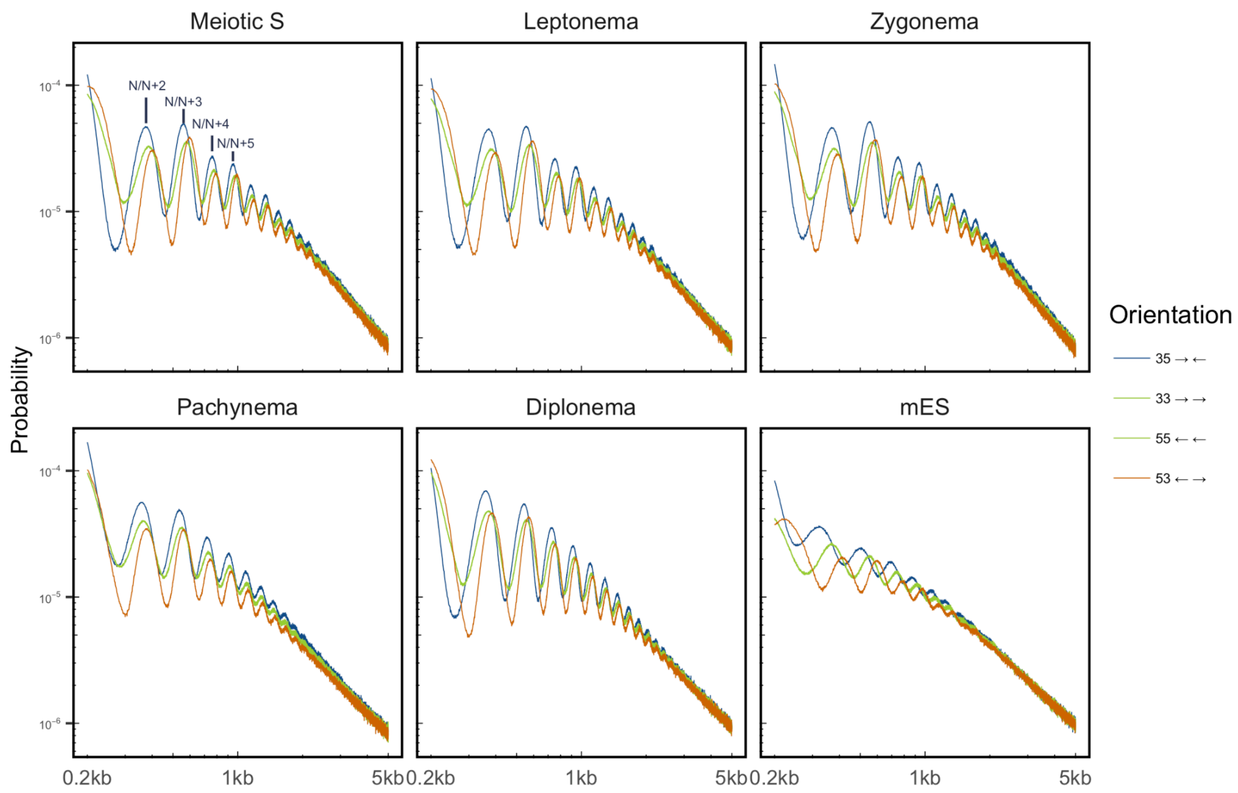


**Figure S3. Contact probability decay at single nucleotide resolution within 5 Kb**

The read pairs at different distances were counted at a resolution of 1 bp. The contact probabilities were calculated by dividing the pair number at each distance by total pairs.The reads were shifted to the nucleosome dyad by adding 73 bp to the forward reads and subtracting 73 bp from the reverse reads.

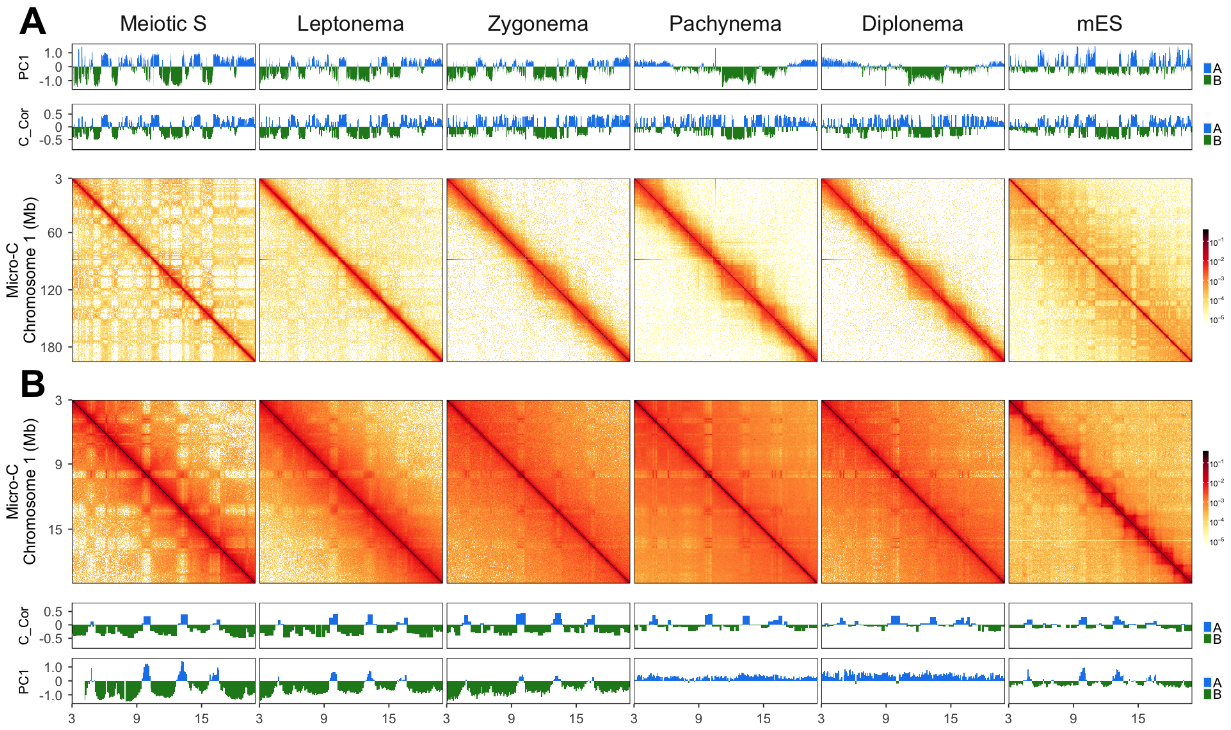


**Figure S4. Genome-wide chromatin reorganization derived from Micro-C**

(A) Top: The profiles of PC1 calculated at a resolution of 100 Kb. Middle: The profiles of correlation scores calculated by Calder (Y. Liu et al. 2021) at a resolution of 50 Kb. Bottom: Matrices of chromosome 1 at a resolution of 100 Kb, plotted from Micro-C data.

(B) A snapshot of Micro-C matrices at chromosome 1: 3 -20 Mb. Top: Matrices were plotted at a resolution of 50 Kb from Micro-C data. Middle: The profiles of correlation scores were generated by Calder at a resolution of 50 Kb, aligned with the regions depicted in the Micro-C matrices. Bottom: The profiles of PC1 were calculated at a resolution of 100 Kb and aligned with the regions depicted in the Micro-C matrices.

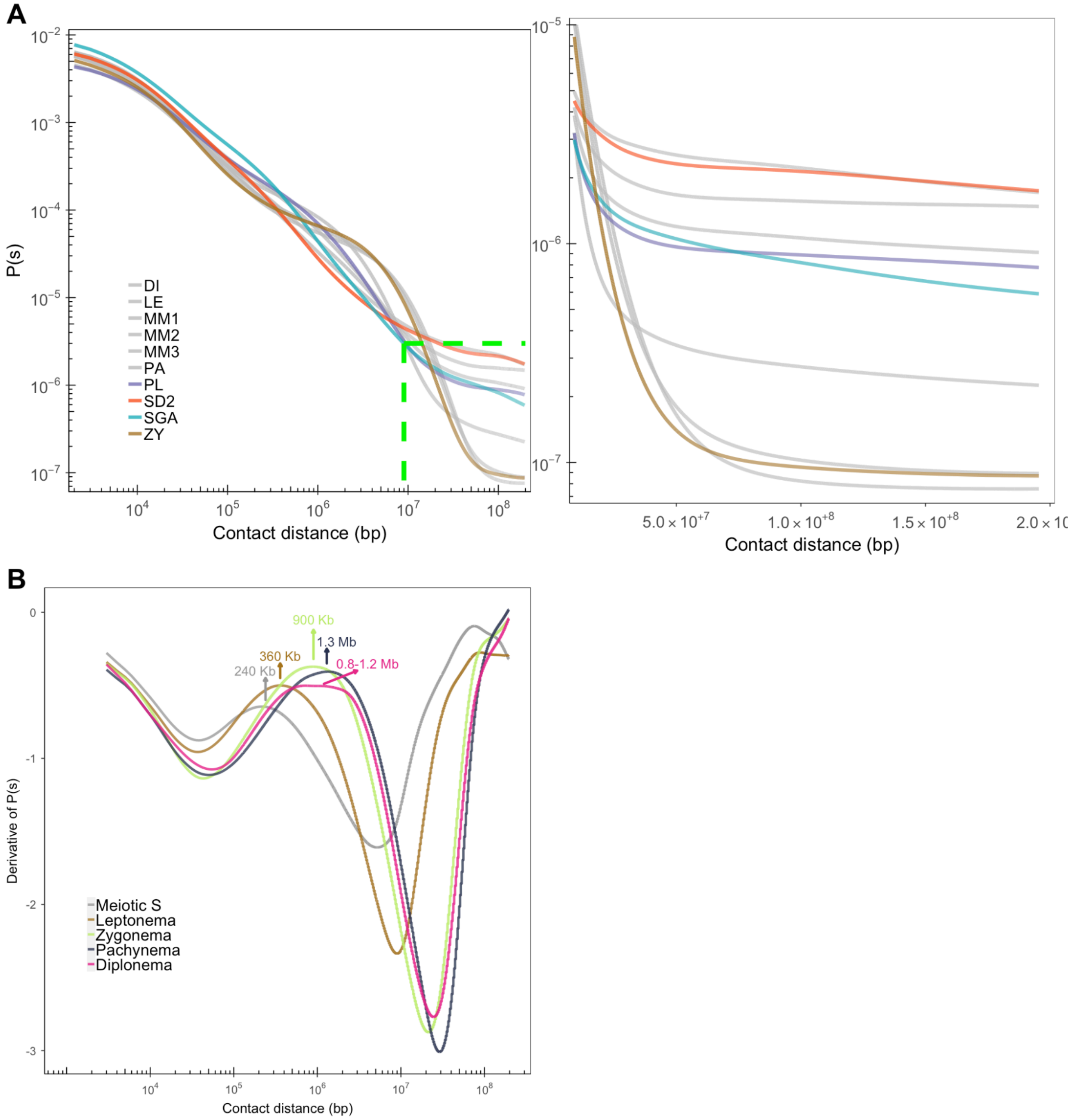


**Figure S5. Contact probability plotted as a function of contact distances and their corresponding derivatives**

(A) Left: The contact probability was calculated at a resolution of 1 Kb. Stages from unDiff.SGA to Meiotic S are indicated by gray color. Stages during MPI are indicated by different colors. The x- and y-axis were plotted at log10 scale. Green dash lines frame the region where far-cis contacts began to diminish. Right: The framed region was plotted separately. In this plot, the x-axis was plotted as a linear scale, the y-axis was plotted as a log10 scale.

(B) The derivatives of P(s) from meiotic S to diplotene stages. The local maxima are indicated as arrow bars, with the estimated loop sizes labeled above the arrows.

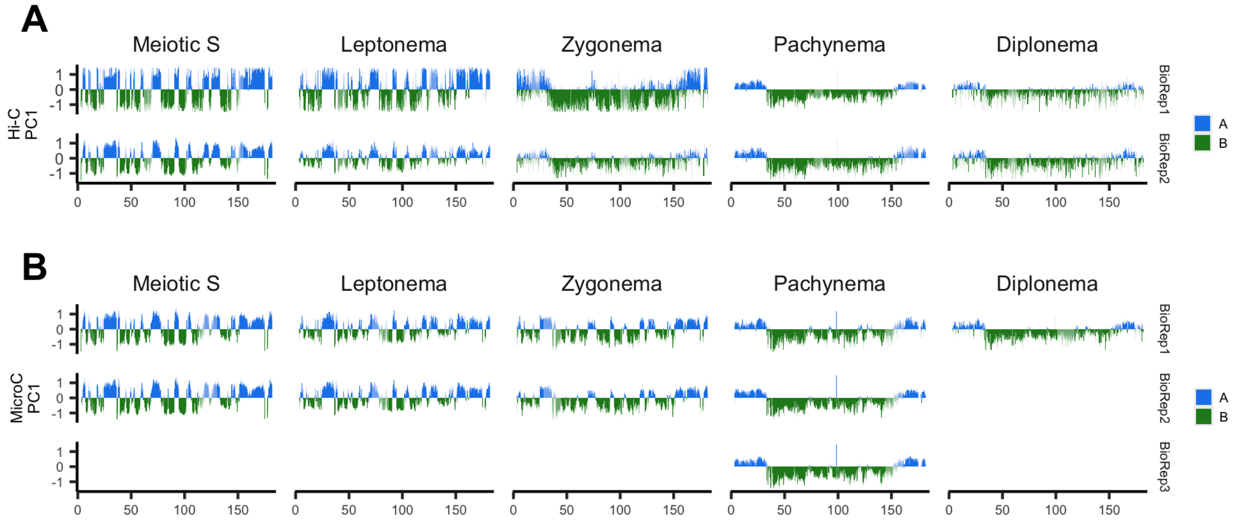


#### Figure S6. PC1 profiles derived from biological replicates of both Micro-C and Hi-C data

(A) PC1 along chromosome 2 for each replicate of Hi-C data. Principal component analysis was performed at a resolution of 100 Kb.

(B) PC1 along chromosome 2 for each replicate of Micro-C data. Principal component analysis was performed at a resolution of 100 Kb.

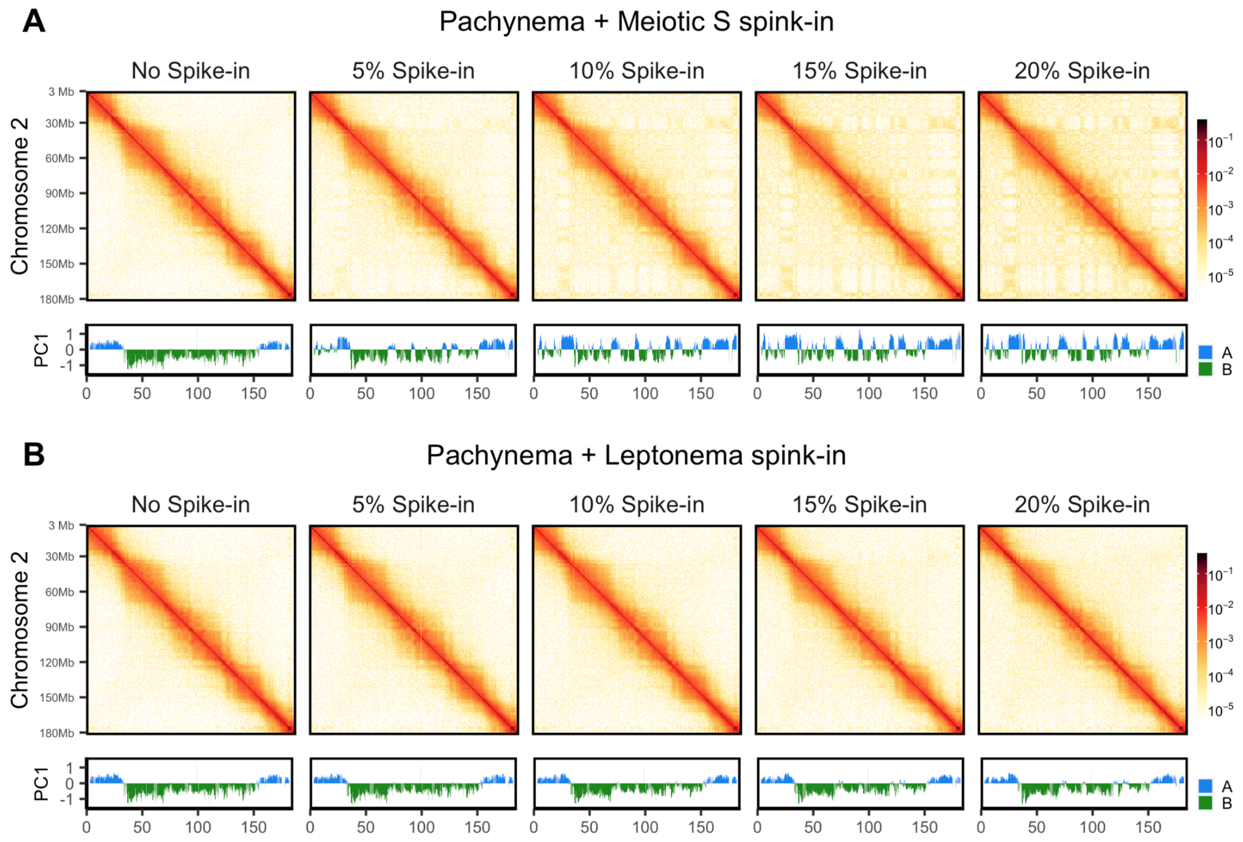


**Figure S7. Identification of compartmentalization by Spike-in.**

(A) Compartmentalization was identified by spiking in interactions from the meiotic S stage. Various percentages of read pairs from the meiotic S stage were integrated with read pairs from the pachytene stage. Subsequently, matrices with a resolution of 100 Kb were constructed and balanced. The matrices for chromosome 2, which include different percentages of spike-in, were then plotted. Additionally, the first principal component (PC1) was calculated from the balanced matrices.

(B) Compartmentalization was identified by spiking in interactions from the leptotene stage. Various percentages of read pairs from the leptotene stage were integrated with read pairs from the pachytene stage. Subsequently, matrices with a resolution of 100 Kb were constructed and balanced. The matrices for chromosome 2, which include different percentages of spike-in, were then plotted. Additionally, the first principal component (PC1) was calculated from the balanced matrices.

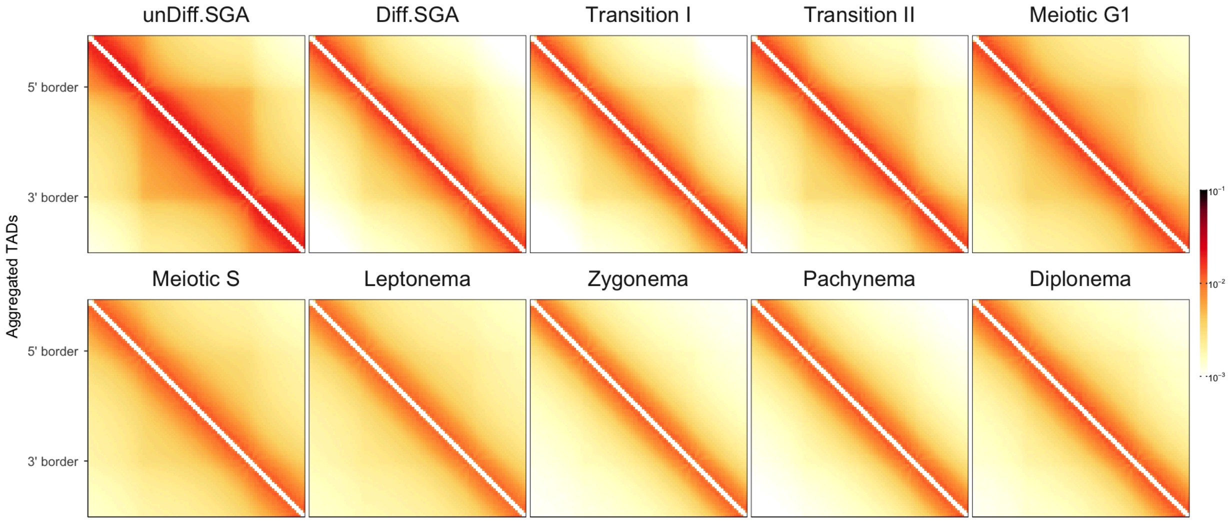


**Figure S8 Aggregated interactions of conserved TADs**

The TAD boundaries were identified from both unDiff.SGA and mouse ES cells (Yan et al. 2018) using Cooltools at a resolution of 10 Kb with a window size of 100 Kb. In order to pinpoint conserved boundaries, we initially extended 10 Kb on both sides of the boundaries identified from unDiff.SGA. Boundaries identified in unDiff.SGA that overlapped with boundaries identified in mES cells were classified as conserved boundaries. TAD domains in unDiff.SGA were defined as regions bordered by the two nearest boundaries. Only the TADs in unDiff.SGA defined by the conserved boundaries were used for this analysis.

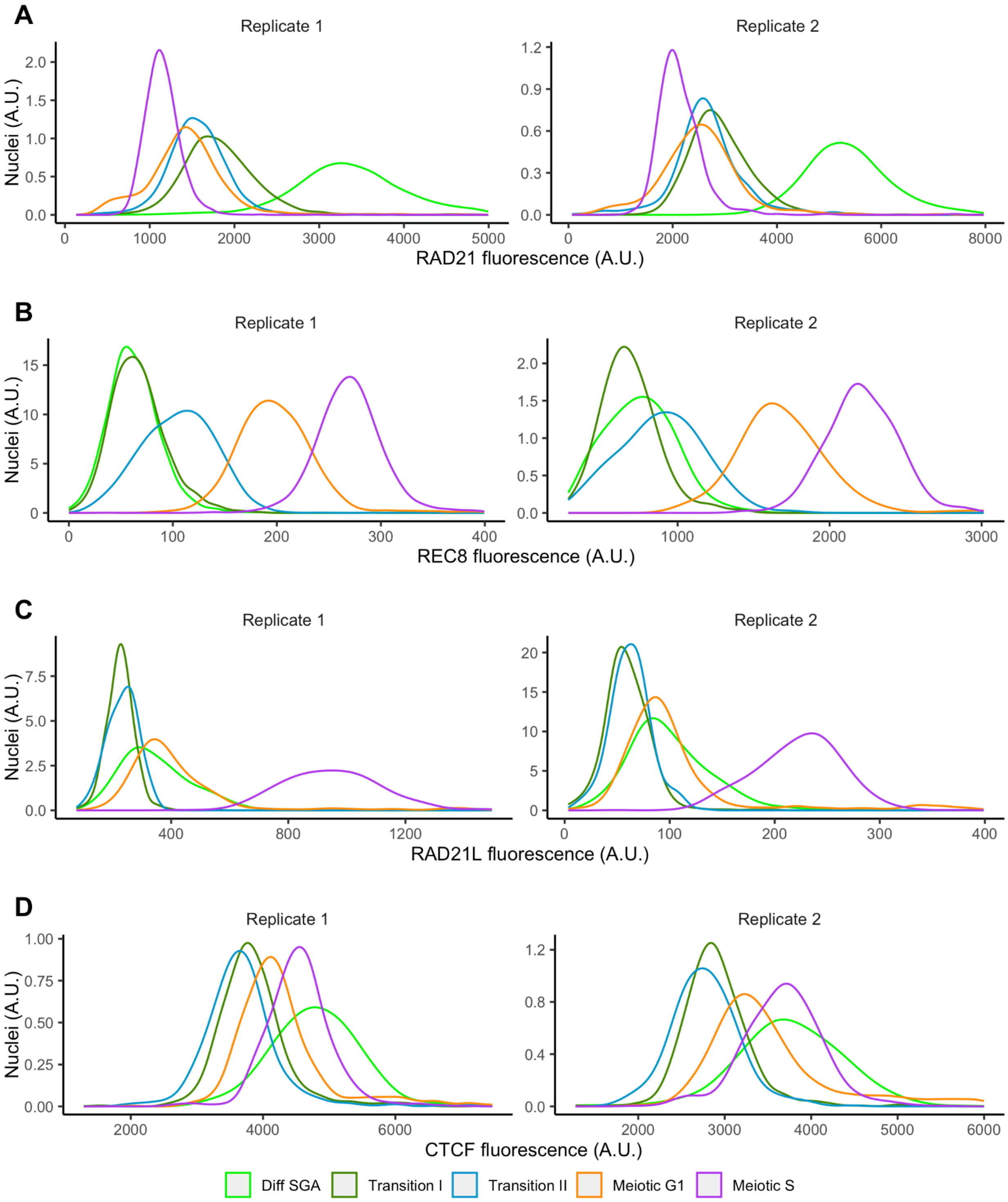

A
Replicate 1
Replicate 2
Nuclei (A.U.)
RAD21 fluorescence (A.U.)
B
Replicate 1
Replicate 2
Nuclei (A.U.)
REC8 fluorescence (A.U.)
C
Replicate 1
Replicate 2
Nuclei (A.U.)
RAD21L fluorescence (A.U.)
D
Replicate 1
Replicate 2
Nuclei (A.U.)
CTCF fluorescence (A.U.)
Diff SGA
Transition I
Transition II
Meiotic G1
Meiotic S

#### Figure S9. Changes in factors that regulate chromatin folding at the scale of TAD

(A) Changes in RAD21. The populations ranging from Diff.SGA to meiotic G1 were gated as shown in Figure S1. The meiotic S population was gated based on a 2-4C DNA content, very low DMRT1 expression, and very strong STRA8 expression. The profiles of RAD21 immunostaining signals from the different gated stages were plotted for two biological replicates.

(B) Changes in REC8. The populations were gated in the same manner as shown in (A). The profiles of REC8 immunostaining signals from the different gated stages were plotted for two biological replicates.

(C) Changes in RAD21L. The populations were gated in the same manner as shown in (A). The profiles of RAD21L immunostaining signals from the different gated stages were plotted for two biological replicates.

(D) Changes in CTCF. The populations were gated in the same manner as shown in (A). The profiles of CTCF immunostaining signals from the different gated stages were plotted for two biological replicates.

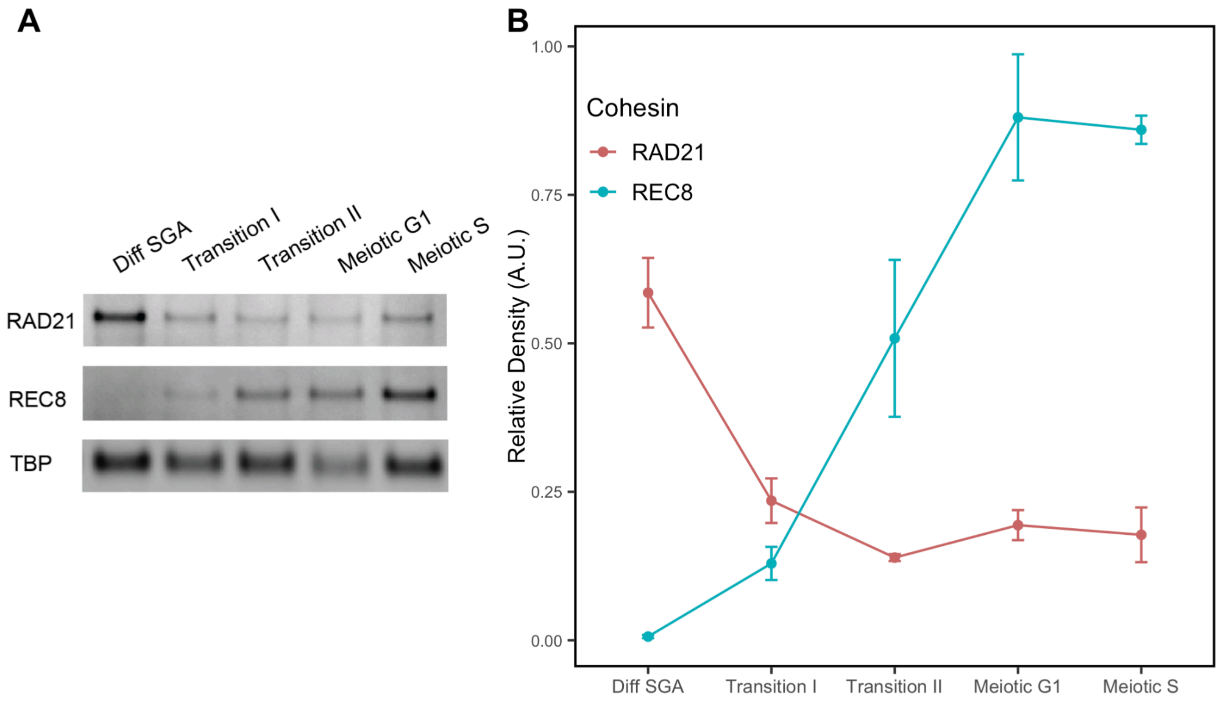


**Figure S10. Western blot analysis of REC8 and RAD21.**

(A) Western blot analysis of mitotic kleisin RAD21 and meiotic kleisin REC8 in different "pre-meiotic" stages. TBP (TATA box binding protein) was used as the loading control.

(B) Quantification of the western blot. Protein abundance at each stage was normalized to the levels of TBP.

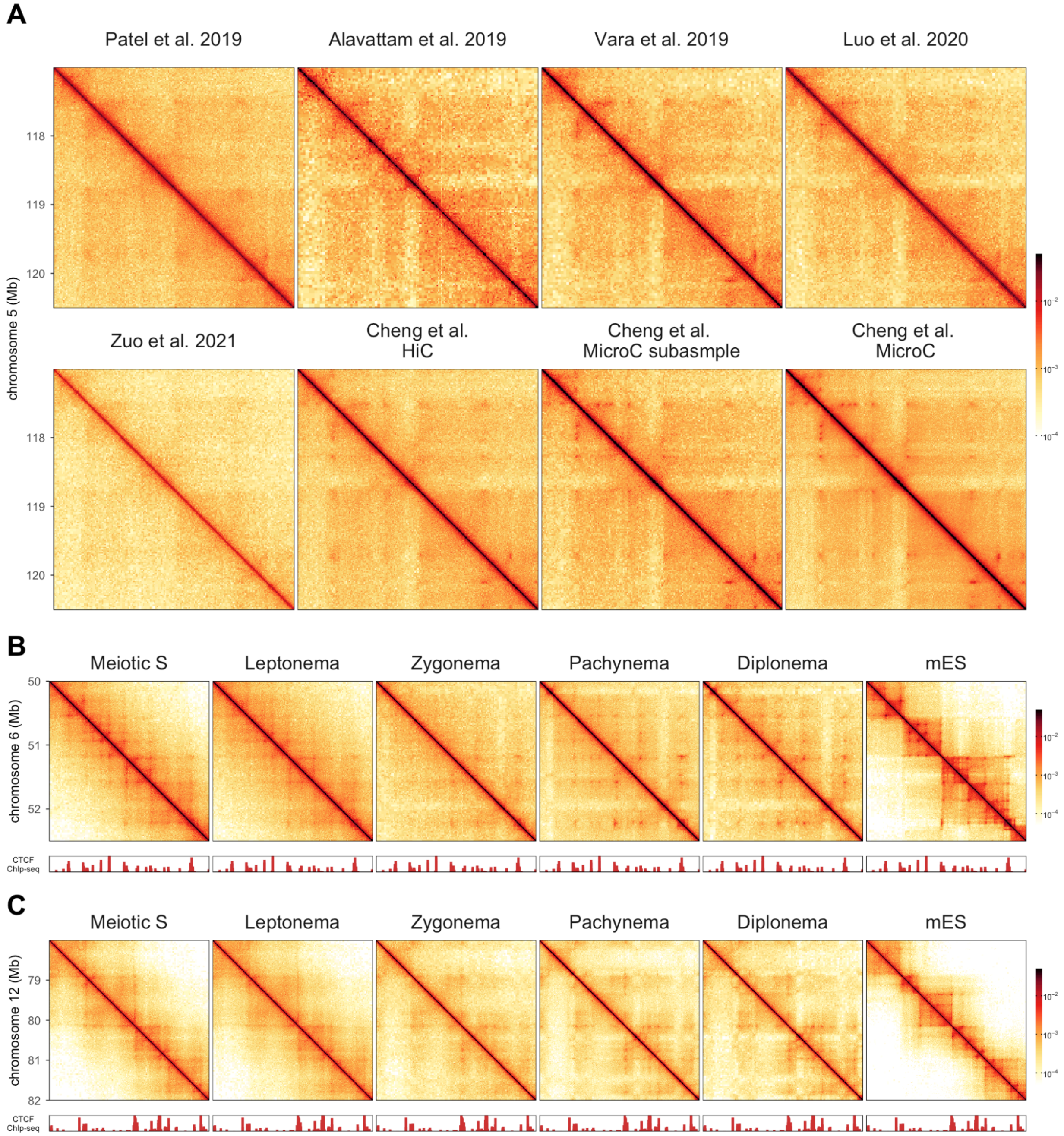


**Figure S11. Dots visualization in Hi-C/Micro-C matrices generated from ours and published data.**

(A) Pachytene matrices were plotted from previously published studies as well as our Hi-C/Micro-C data. The matrices show a region spanning from 117 Mb to 120 Mb on chromosome 5.

Patel et al. 2019: Data from Francesca Cole and Kevin D. Corbett's group. Total read pairs: 478 million;

Alavattam et al. 2019: Data from Satoshi H Namekawa's group. Total read pairs: 284 million;

Vara et al. 2019: Data from Aurora Ruiz-Herrera's group. Total read pairs: 411 million;

Luo et al. 2020: Data from Xiaoyuan Song's group. Total read pairs: 205 million;

Zuo et al. 2021: Data from Ming Lei and Qian Bian's group. Total read pairs:336 million;

Cheng et al. HiC: Data from this study. Total read pairs: 634 million;

Cheng et al. Micro-C subsample: Data from this study. The total valid read pairs were subsampled to 690 million;

Cheng et al. Micro-C: Data from this study. Total read pairs: 2.92 Billion

(B) A snapshot of the Micro-C matrix of chromosome 6 at a resolution of 5 Kb (region: 50 Mb to 52.5 Mb). CTCF ChIP coverage is plotted at the bottom;

(C) A snapshot of the Micro-C matrix of chromosome 12 at a resolution of 5 Kb (region: 78 Mb to 82 Mb). CTCF ChIP coverage is plotted at the bottom;

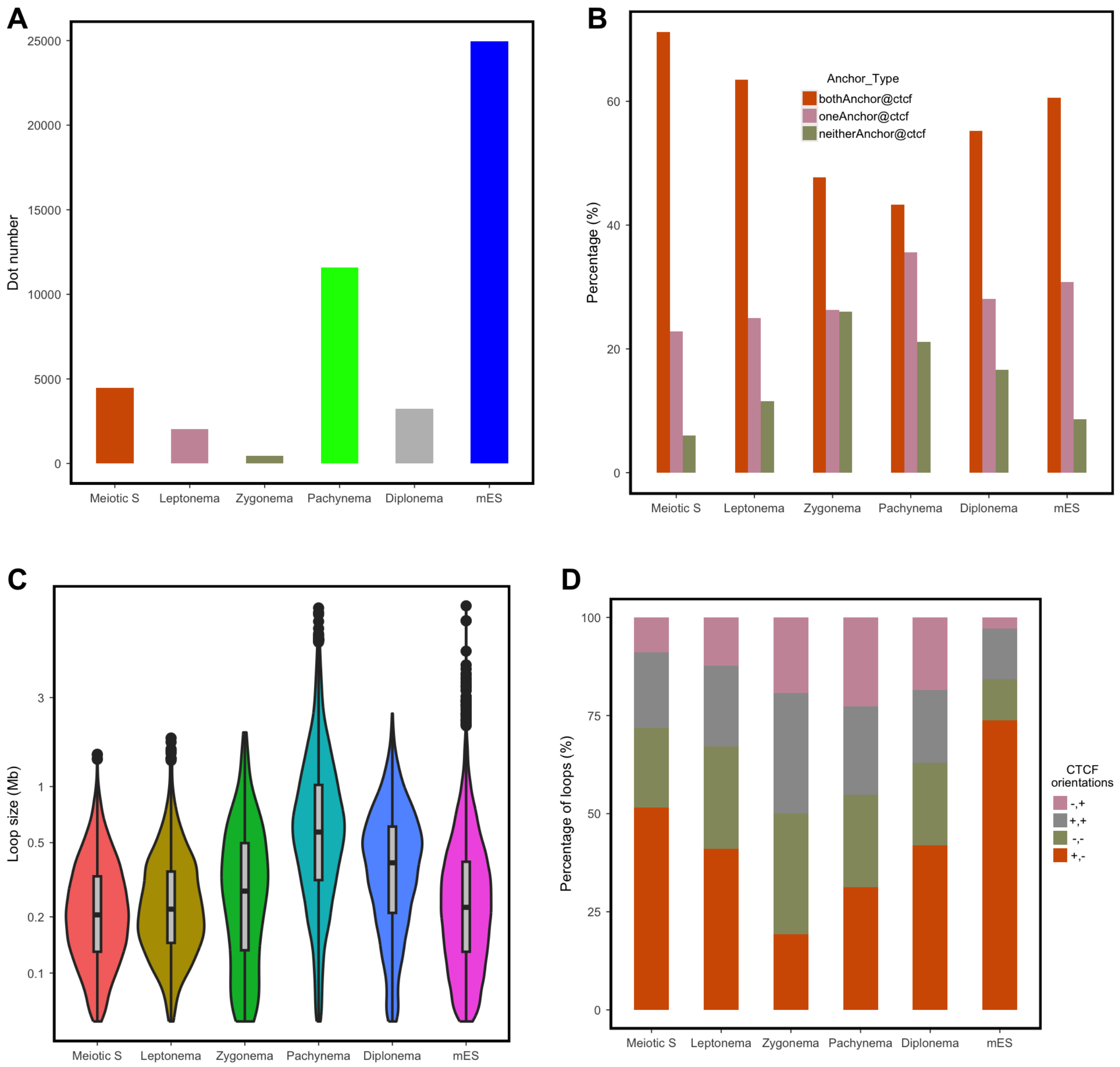


**Figure S12. A summary of all dots identified from each stage**

(A) The number of dots identified from each cell type. The anchors that are less than 50 Kb were removed because the calling at this range was not accurate enough;

(B) The percentage of dots with both anchors, one anchor, and no anchor overlapping with CTCF binding sites. CTCF motifs were identified from CTCF chip-seq peaks by FIMO. The motifs were extended +/- 5 kb when overlapping the anchors;

(C) Distance between anchors of the identified dots;

(D) Percentage of dots with different oriented anchors. Anchors that overlap with a single CTCF motif were selected for this analysis.

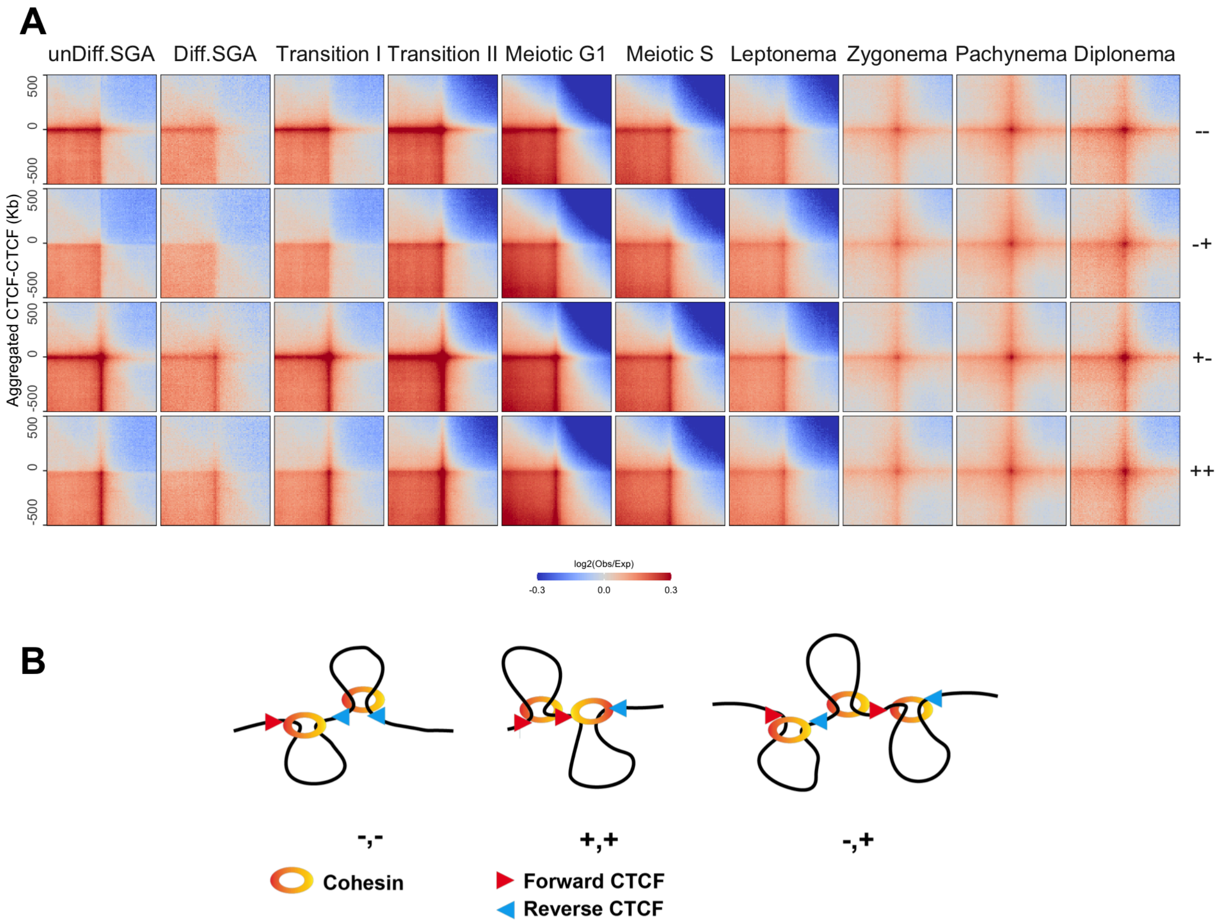


**Figure S13. Aggregated interactions of pairwise CTCF from different orientations.**

(A) Aggregated interactions between CTCF of different orientations. Each map is plotted at a resolution of 10 Kb, with a flanking region of 500 Kb. The bottom row is the same plot shown in Figure 2C.

(B) A schematic to show how the Hi-C loops are formed between non-convergent CTCFs. This schematic is adapted from (N. Q. Liu et al. 2021)

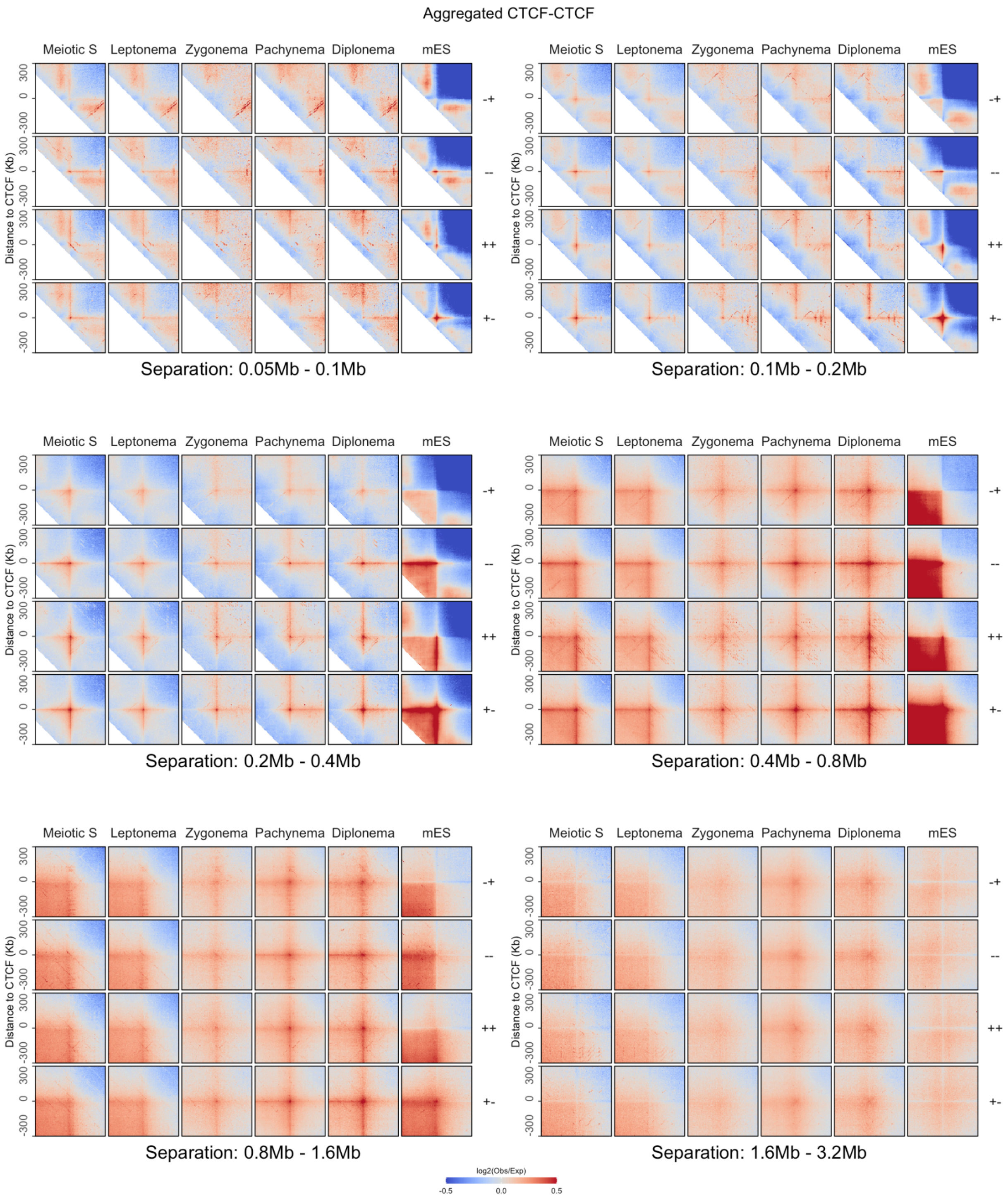


**Figure S14. Aggregation of pairwise CTCF binding sites categorized by orientation and distance.**

Aggregated interactions between CTCF binding sites was plotted at a resolution of 10 Kb, with a flanking region of 300 Kb. The orientation was labeled at the right of each panel. +: CTCF binding motifs with forward orientation; -: CTCF binding motifs with reverse orientation. The separation between two CTCF binding motifs was labeled at the bottom of each panel.

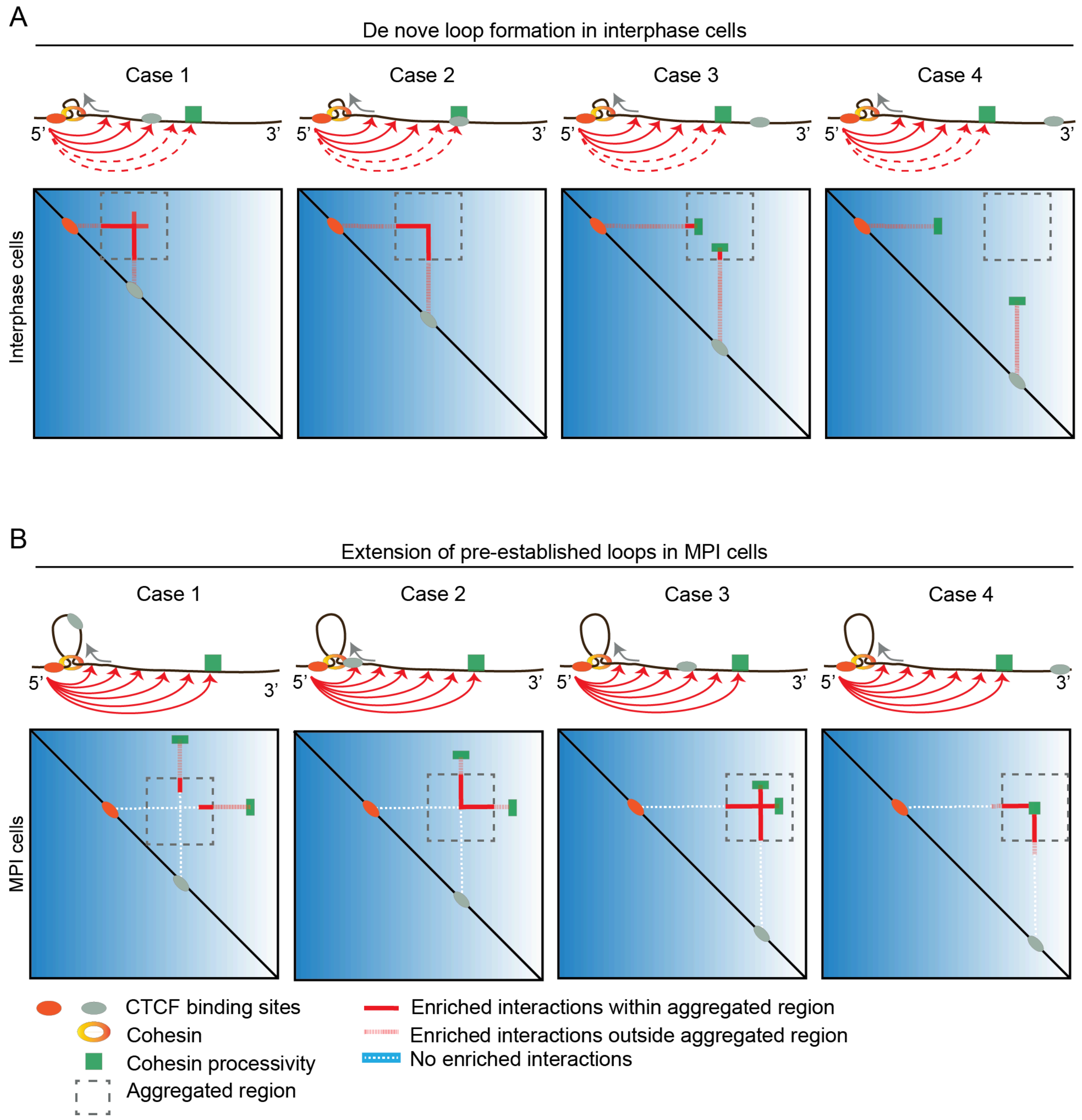


**Figure S15. The schematic illustrates the enriched interaction pattern at various distances between CTCF binding sites and cohesin processivity.**

(A) In somatic interphase cells, cohesin is loaded to chromosomes and generates loops from the beginning.

(B) In MPI cells, cohesins are pre-loaded. The loops are extended from preformed loops.

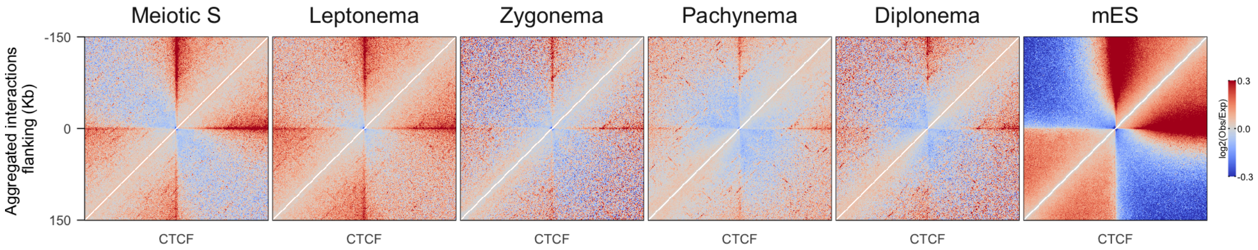


**Figure S16. A zoomed-in view of Figure 4F illustrates the depletion of short-range interaction enrichment.**

Aggregation of interactions surrounding CTCF binding sites with unified motif orientation. The analysis was performed at a resolution of 1 Kb, with a 150 Kb flanking region. Only the interactions at loci with forward-oriented CTCF was shown.

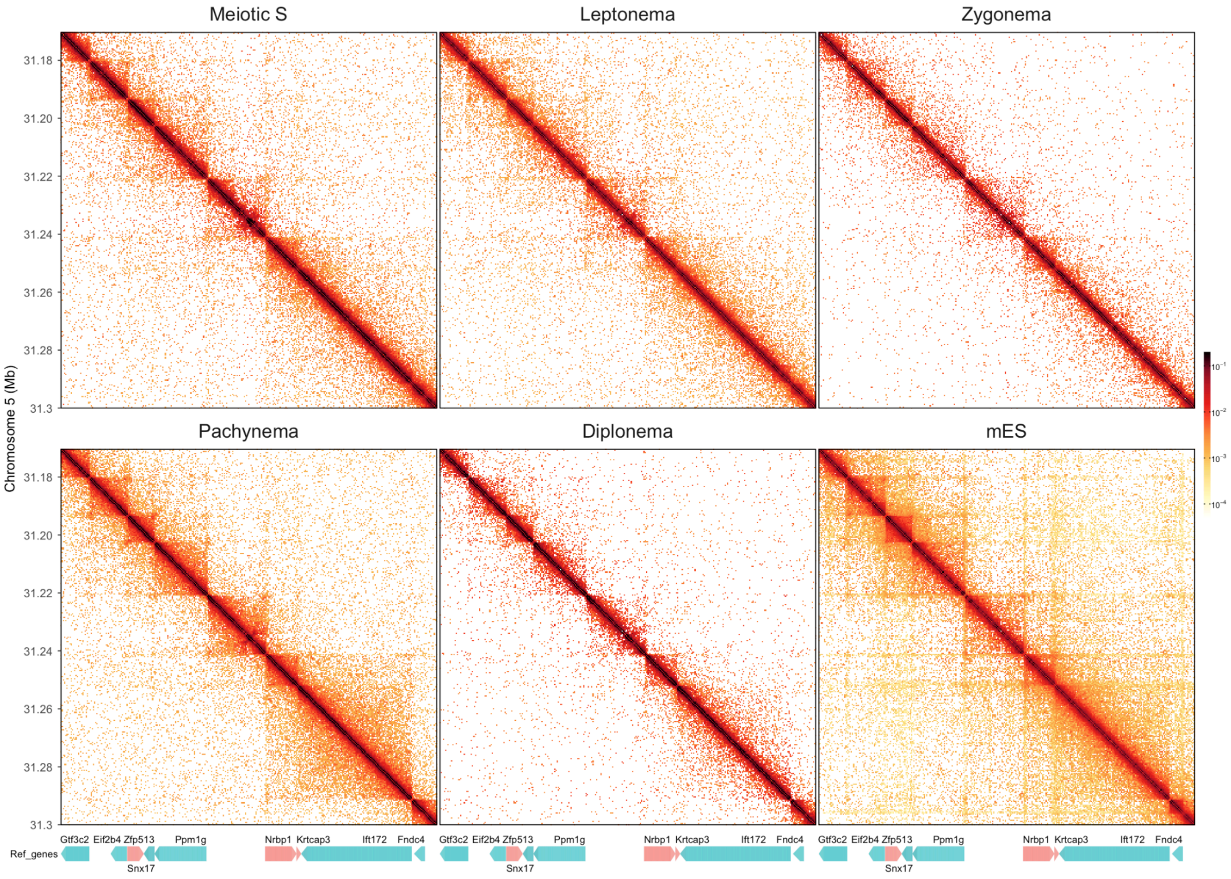


**Figure S17. Snapshots of Micro-C matrices of at a resolution of 400 bp**

Micro-C matrices from meiotic S through MPI were plotted for the region spanning 31 Mb to 31.4 Mb on chromosome 5 at a resolution of 400 bp. The Micro-C matrix of the same region from mES(Hsieh et al. 2020) was plotted as a comparison. Genes in the same region were annotated at the bottom. Bars with arrows indicate the orientation of the genes. The difference of visibility mainly results from sequencing depth.

Meiotic S: 1371 million read pairs

Leptonema: 1348 million read pairs

Zygonema: 752 million read pairs

Pachynema: 3263 million read pairs

Diplonema: 996 million read pairs

mES: 3175 million read pairs

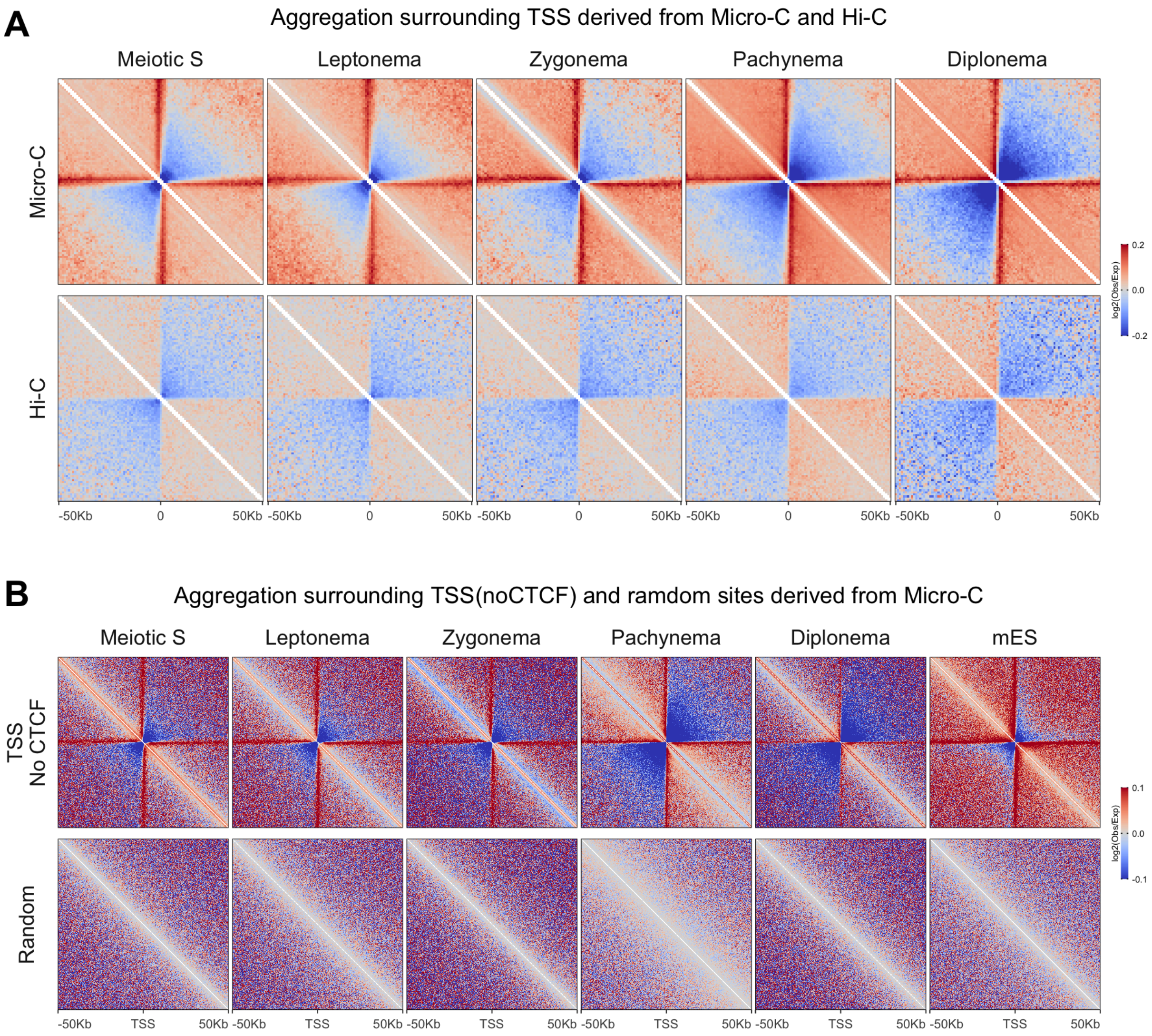


#### Figure S18. Aggregated interactions surrounding TSSs.

(A) Aggregated interactions centered around TSSs were calculated for both Micro-C and Hi-C data. TSSs were derived from USCS annotated genes. The aggregated interactions were calculated at a resolution of 1000 bp with a flanking region of 50 Kb.

(B) Aggregated interactions centered around TSSs and random sites were calculated for Micro-C data. TSSs were derived from USCS annotated genes. Any TSS with CTCF binding sites within ± 5 Kb were excluded from the analysis. The aggregated interactions were calculated at a resolution of 200 bp with a flanking region of 50 Kb.

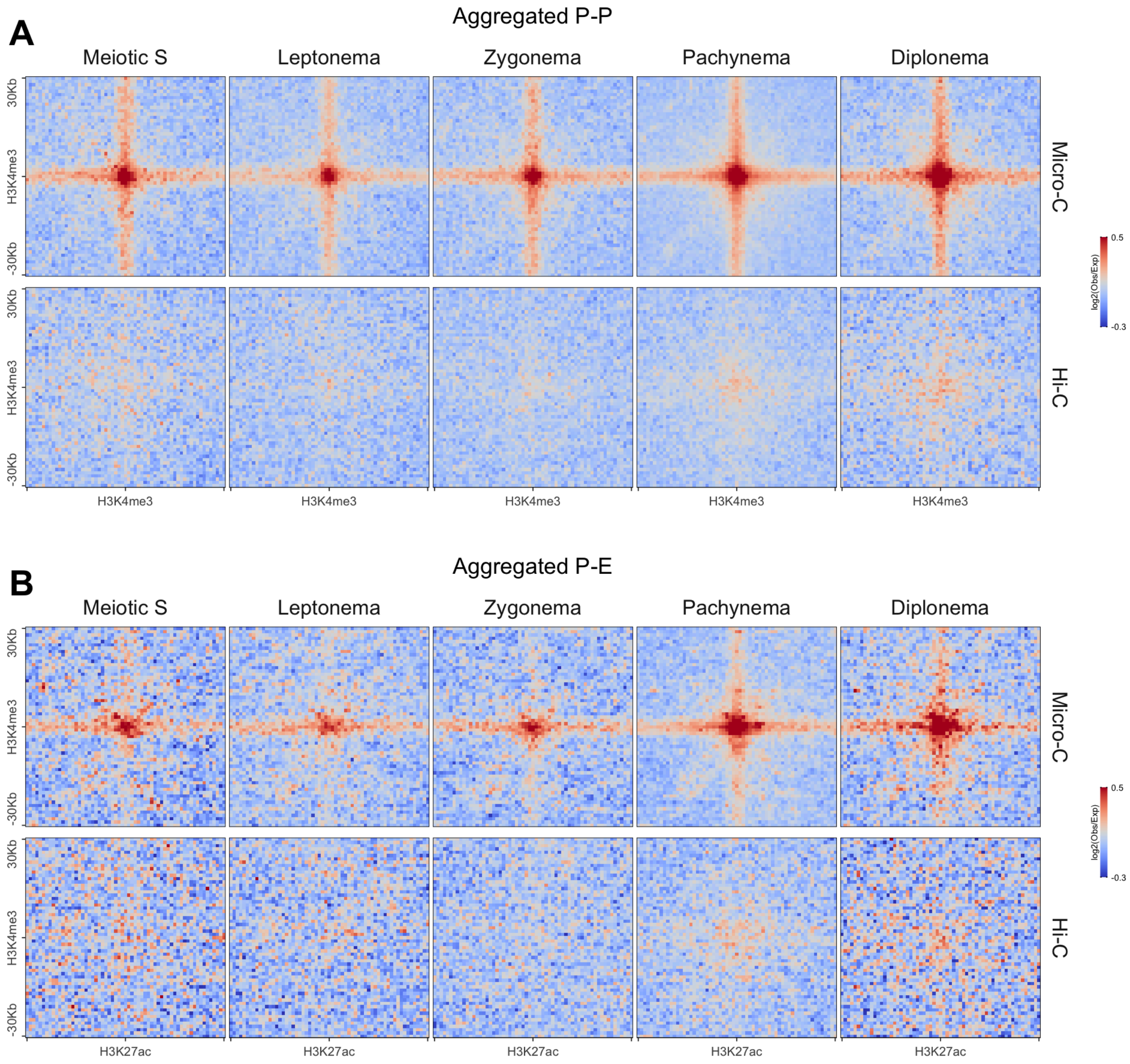


#### Figure S19. Aggregated interactions between regulatory elements

(A) Aggregated promoter-promoter interactions were plotted for both Micro-C and Hi-C data. The promoters were derived from H3K4me3 Chip-seq data (Lam et al. 2019). The H3K4me3 modification sites that overlapped with hotspots were excluded from calculations. Two promoters at distances ranging from 5 Kb to 5 Mb were paired and aggregated at a resolution of 1000 bp with a flanking region of 50 Kb.

(B) Aggregated promoter-enhancer interactions were plotted for both Micro-C and Hi-C data. The enhancers were derived from H3K27ac Chip-seq data (Lam et al. 2019 ;Maezawa et al. 2020). Common peaks from both studies were selected and only the sites that did not overlap with H3K4me3 peaks and hotspots were used for the calculation. Promoters and enhancers at distances ranging from 5 Kb to 5 Mb were paired and aggregated at a resolution of 1000 bp with a flanking region of 50 Kb.

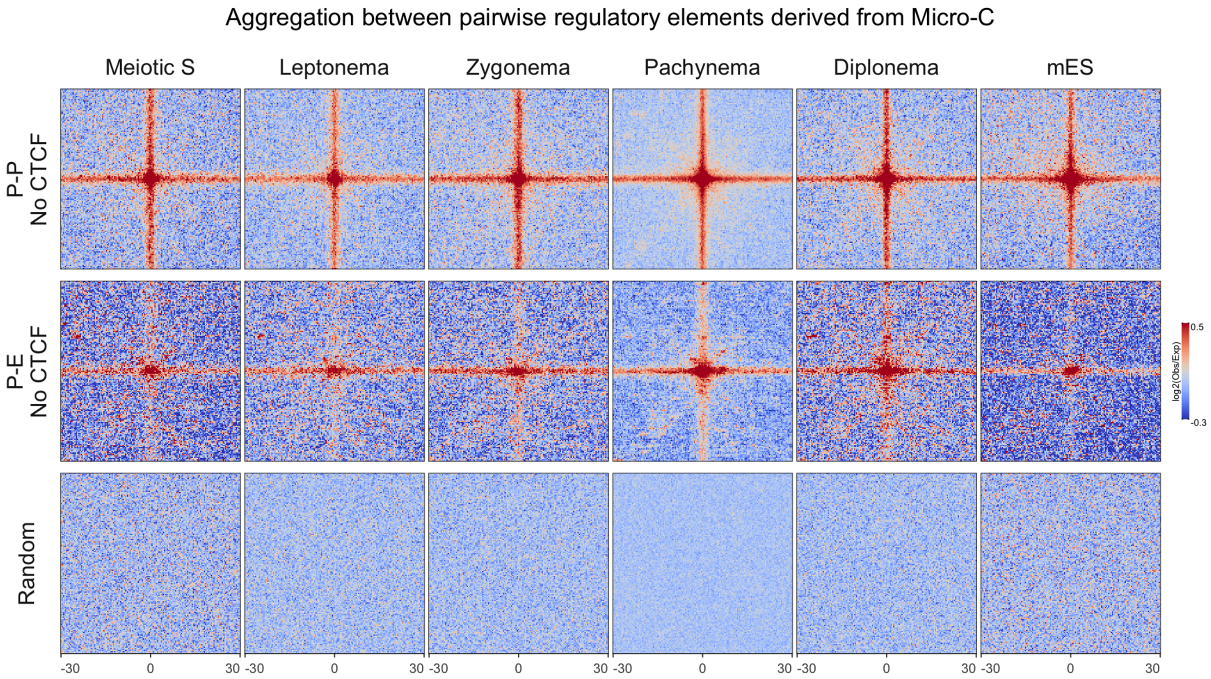


#### Figure S20. Aggregation of P-P/P-E interactions that do not overlap with CTCF

Similar to Figure 5 and Figure S19, promoters were identified based on H3K4me3 ChIP-seq data (Lam et al. 2019). Enhancers were defined as the common peaks found in two separate H3K27ac ChIP-seq datasets (Lam et al.2019; Maezawa et al. 2020). Peaks from these datasets that do not overlap with H3K4me3 peaks were considered as enhancers. Furthermore, both promoters and enhancers that coincided with CTCF binding sites within a range of ± 5 Kb were excluded from the calculations. Two promoters, or promoters and enhancers, at distances ranging from 5 Kb to 5 Mb were paired and aggregated at a resolution of 1000 bp with a flanking region of 50 Kb.

A

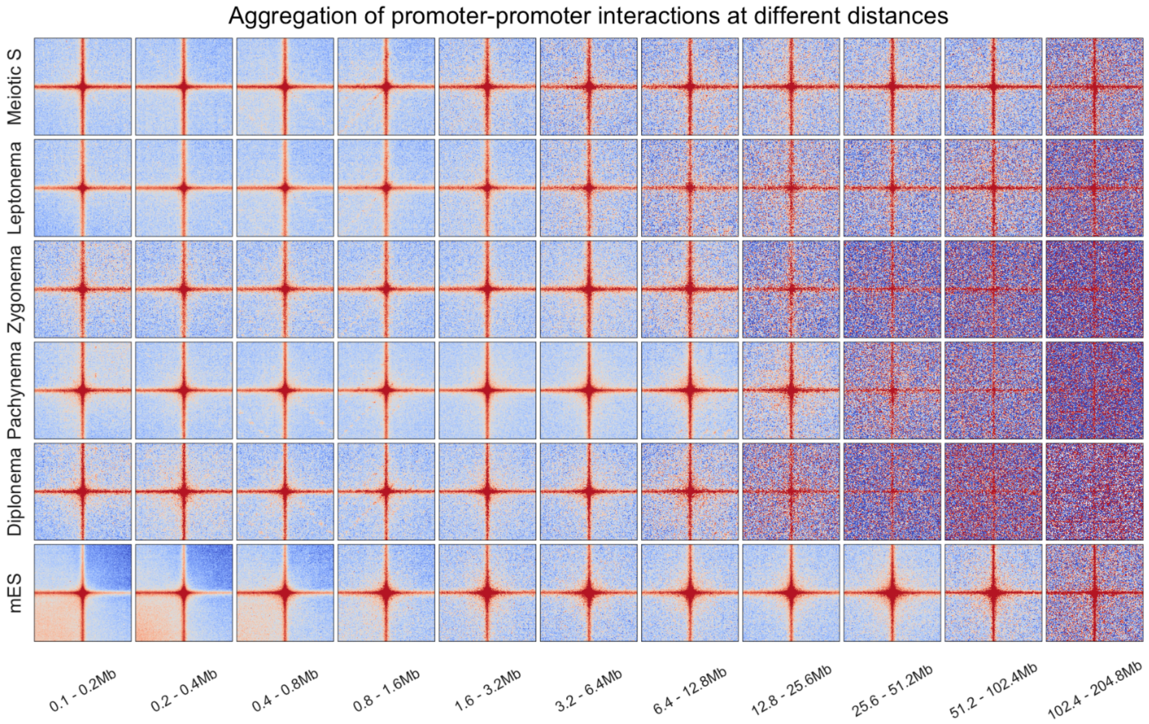
Aggregation of promoter-promoter interactions at different distances
Meiotic S
Leptonema
Zygonema
Pachynema
Diplonema
mES
0.1 - 0.2Mb
0.2 - 0.4Mb
0.4 - 0.8Mb
0.8 - 1.6Mb
1.6 - 3.2Mb
3.2 - 6.4Mb
6.4 - 12.8Mb
12.8 - 25.6Mb
25.6 - 51.2Mb
51.2 - 102.4Mb
102.4 - 204.8Mb

B

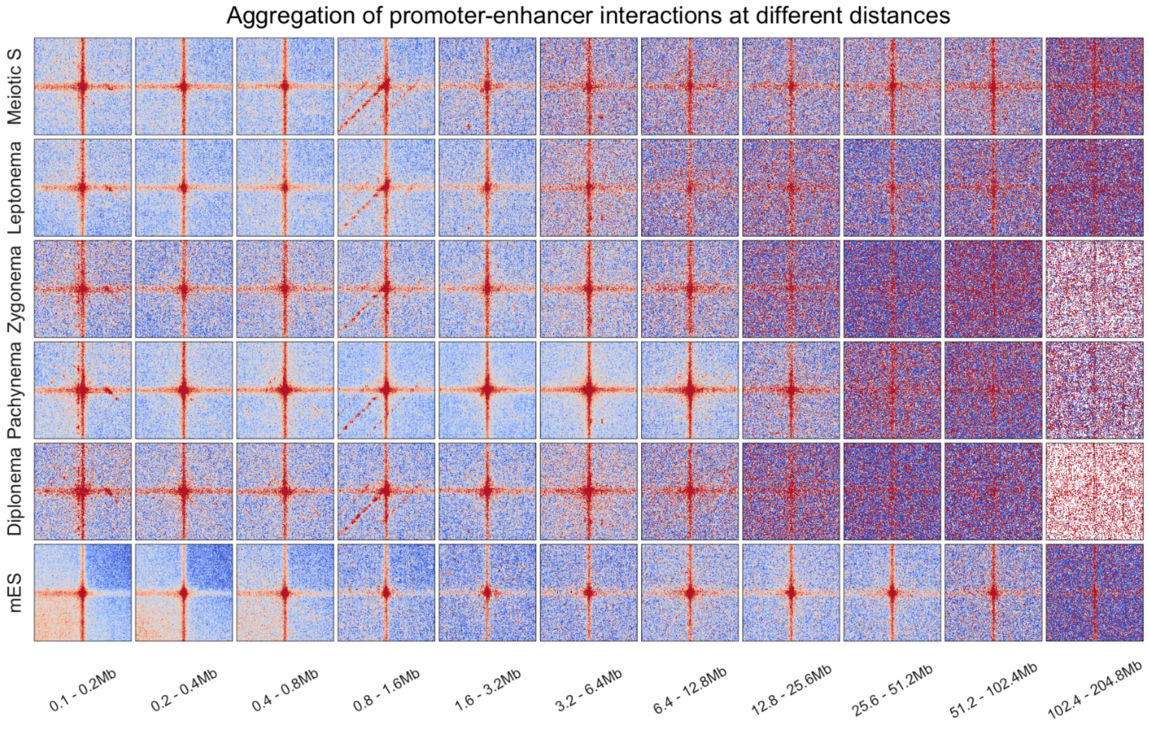
Aggregation of promoter-enhancer interactions at different distances
Meiotic S
Leptonema
Zygonema
Pachynema
Diplonema
mES
0.1 - 0.2Mb
0.2 - 0.4Mb
0.4 - 0.8Mb
0.8 - 1.6Mb
1.6 - 3.2Mb
3.2 - 6.4Mb
6.4 - 12.8Mb
12.8 - 25.6Mb
25.6 - 51.2Mb
51.2 - 102.4Mb
102.4 - 204.8Mb

##### Figure S21. Aggregation of P-P/P-E interactions at different distances

(A) Aggregated promoter-promoter interactions at different distances. The same promoters dataset in Figure 5 were used here. Paired promoters were categorized based on distances and aggregated at a resolution of 400 bp with a flanking region of 30 Kb.

(D) Aggregated promoter-enhancer interactions at different distances. The same promoters and enhancer dataset in Figure 5 were used here. Paired promoter-enhancers were categorized based on distances and aggregated at a resolution of 400 bp with a flanking region of 30 Kb.

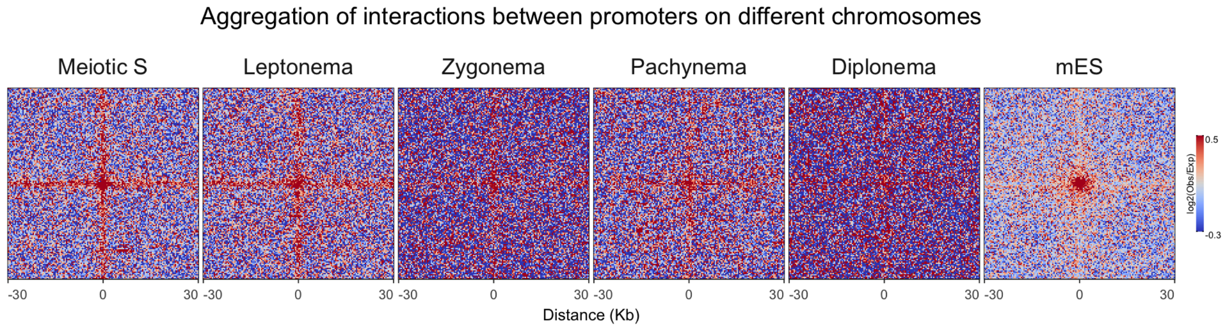


#### Figure S22. Aggregation of interactions at promoters from different chromosomes

The same promoters dataset in Figure 5 were used here. All promoters from different chromosomes were paired and aggregated at a resolution of 400 bp with a flanking region of 30 Kb.
